# Assembly, Annotation, and Comparative Analysis of the Mitochondrial Genome of Winged Bean (*Psophocarpus tetragonolobus*): Insights into Evolutionary Adaptation and Codon Usage Bias

**DOI:** 10.1101/2025.01.24.634632

**Authors:** Nikhil Kumar Singh, Binay K. Singh, Piyush Kumar, Avinash Pandey, Sudhir Kumar, Sujit Kumar Bishi, A. Pattanayak, V.P. Bhadana, Sujay Rakshit, Kishor U. Tribhuvan

**Author notes:** ICAR-Indian Institute of Wheat and Barley Research, Karnal 132001, Haryana, India.

## Abstract

The winged bean (*Psophocarpus tetragonolobus*) is an underutilized, nutritionally dense legume with significant agricultural potential due to its adaptability to tropical environments and nitrogen-fixing capabilities. This study presents the assembly, annotation, and comparative analysis of its complete mitochondrial genome, spanning 366,925 bp. The genome comprises 64 genes, including 38 protein-coding genes, 12 tRNA genes, and 6 rRNA genes. Codon usage analysis revealed moderate bias in mitochondrial genes, primarily shaped by mutational pressures, while tRNA gene content aligns with codon preferences, optimizing translational efficiency. Phylogenetic analysis placed *P. tetragonolobus* within the Phaseoleae tribe, clustering closely with *Glycine max* and *Vigna radiata*, while synteny analysis highlighted high genomic conservation with related legumes. Ka/Ks ratio analysis underscored strong purifying selection in core mitochondrial genes, reflecting functional constraints essential for energy metabolism. Comparative analyses of mitochondrial and chloroplast genomes revealed distinct evolutionary pressures, with chloroplast genes exhibiting more significant variability influenced by functional roles in photosynthesis. These findings provide valuable insights into the evolution and adaptation of *P. tetragonolobus*, establishing a foundation for its potential application in sustainable agriculture and crop improvement programs.

## Introduction

The study of plant mitochondrial genomes, with their unique and complex characteristics, holds substantial importance due to their roles in energy production, cellular signalling, and responses to environmental stress (Douce 2012)). The significant contribution of mitochondria in cell functioning is attributed to its genetic system, which is semi-autonomous (Douce 2012; Day et al. 2013). Plant mitochondrial genomes exhibit unique characteristics, including structural complexity, recombination events, and horizontal gene transfer, distinguishing them from animal mitochondria (Bock & Knoop 2012; Mower et al. 2012). The large genome size and complexity are partially attributed to accumulating non-coding sequences, duplications, and foreign DNA sequences, such as those originating from nuclear and chloroplast genomes (Gualberto & Newton 2017). While challenging for genome assembly and annotation, these variabilities provide critical insights into the unique mechanisms driving plant genomic evolution and structural rearrangements (Tomohiko Kubo & Newton 2008).

Structural diversity within plant mitochondrial genomes is shaped by recombination processes that result in multipartite genome structures, where circular or linear arrangements can form due to repeated sequences (Bock & Knoop 2012)). These repeated sequences enable homologous recombination, generating subgenomic molecules that vary in copy number within the cell, contributing to an overall dynamic mitochondrial genomic structure (Alexandre Maréchal & Brisson 2010). Such recombination is crucial for maintaining the mitochondrial genome’s integrity under stress conditions and is thought to play a role in adaptation by facilitating genetic diversity within organellar populations (Arrieta-Montiel & Mackenzie 2011). Introns, another notable feature of plant mitochondrial genomes, add to this complexity and are involved in gene regulation and splicing mechanisms that can influence mitochondrial gene expression, particularly under fluctuating environmental conditions (Daniell & Chase 2007). Furthermore, plant mitochondrial genomes often harbour sequences transferred from the nuclear and chloroplast genomes, a process is known as horizontal gene transfer (HGT), which can result in functional gene gains and enhance adaptability to various environmental stresses (Adams & Palmer 2003). This gene transfer between organelles and the nuclear genome may provide adaptive advantages in response to biotic and abiotic stressors, contributing to resilience in diverse environments (Gualberto & Newton 2017).

Identifying unique mitochondrial genome features in underexplored legume species can uncover novel genes or regulatory elements that may be applied to crop improvement programs, contributing to increased agricultural productivity and resilience (Kubo & Mikami 2007). The winged bean (*Psophocarpus tetragonolobus*), sometimes called the “supermarket on a stalk,” is a versatile and nutrient-rich legume but remains scientifically unexplored. This plant, native to Southeast Asia, Africa, and the Pacific Islands, is highly prized for being completely edible, with all its parts—including leaves, tubers, seeds, and flowers—usable for consumption. Known for its high protein content, the winged bean’s seeds contain up to 35% protein, rivalling soybeans and making it an essential dietary protein source in regions where meat and other protein-rich foods may be limited (National Academy of Sciences, 1975). Additionally, the plant’s tubers contain a considerable amount of starch and essential minerals, further contributing to its role as a valuable, multi-purpose crop in areas prone to food insecurity (Rakvong et al. 2024; Murthazar Naim 2020).

Beyond its nutritional benefits, the winged bean plays a vital role in sustainable agriculture. As a member of the legume family, *P. tetragonolobus* has a symbiotic relationship with nitrogen-fixing bacteria, which allows it to enrich the soil with nitrogen—a crucial nutrient that supports crop growth and reduces the need for synthetic fertilisers (Lepcha et al. 2017; Tanzi et al. 2019). This quality makes it a beneficial crop for intercropping systems, particularly in resource-limited and ecologically sensitive areas. Moreover, the winged bean is well-adapted to hot, humid climates and can thrive in poor soils where other crops struggle, making it a resilient option for addressing the challenges posed by climate change (Nwokolo 1996). Given its adaptability, high nutritional value, and ecological benefits, the winged bean has attracted increasing interest in crop diversification and food security programs, especially in regions prone to climate variability.

In recent years, the winged bean has gained attention as a potential’ future crop’ for its resilience and potential health benefits. Its seeds are rich in essential amino acids, vitamins, and antioxidants, which contribute to improved nutrition and immune support for people in tropical regions (Lepcha et al. 2017). Researchers are also exploring the winged bean’s secondary metabolites, which may offer anti-inflammatory and antimicrobial properties, positioning it as a crop with both culinary and medicinal uses (Susanti et al. 2022; Bepary et al. 2023). Due to its protein content, resilience, and diverse applications, the winged bean holds promise for global agricultural sustainability, particularly in the context of rising food demand and the need for environmentally friendly farming practices.

Studying the mitochondrial genome of the winged bean is crucial for several reasons, given its unique nutritional, agricultural, and ecological roles. Mitochondria are essential for cellular respiration and energy production, which directly affects plant growth, productivity, and resilience under stress. In particular, understanding the mitochondrial genome can reveal key insights into how winged bean adapts to challenging environments, such as high temperatures, poor soil conditions, and limited water availability—conditions common in tropical regions where this crop is widely grown (Kumar & Singh 2020; Feng & Xu 2018). By studying the mitochondrial genes associated with energy metabolism, researchers can explore mechanisms that might enhance the crop’s ability to tolerate stress, an important trait for improving yield stability in changing climates (Arrieta-Montiel & Mackenzie 2011).

Lastly, the winged bean’s mitochondrial genome is of evolutionary interest. Plant mitochondrial genomes are highly variable in size and structure across species, with significant rearrangements, repeated sequences, and foreign DNA integrations from nuclear and chloroplast genomes. Studying the winged bean’s mitochondrial genome can provide insights into the evolutionary relationships within the legume family and identify genomic adaptations unique to this species. Such research could illuminate the evolutionary pressures and genetic changes that have enabled winged bean to thrive in tropical conditions, contributing to the broader understanding of plant mitochondrial evolution and adaptation (Kubo & Newton, 2008).

This study aims to assemble and annotate the complete mitochondrial genome of *P. tetragonolobus*, analyze its structural and functional features, and compare its evolutionary traits to related species. We aim to provide valuable insights for future research and crop improvement initiatives by addressing gaps in our understanding of this crop’s mitochondrial genome.

## Results

### Assembly of the Mitochondrial Genome

The complete mitochondrial genome of *P. tetragonolobus* was successfully assembled from filtered high-quality Illumina short reads (523303993 reads) and PacBio long reads (3709181 reads, 63.7 Gbp). The assembled mitochondrial genome spans 366,925 bp (Fig 1), with an average coverage depth of approximately 871X (Supplementary Fig S1), providing high confidence in the base accuracy throughout the genome.

**Figure 1:**
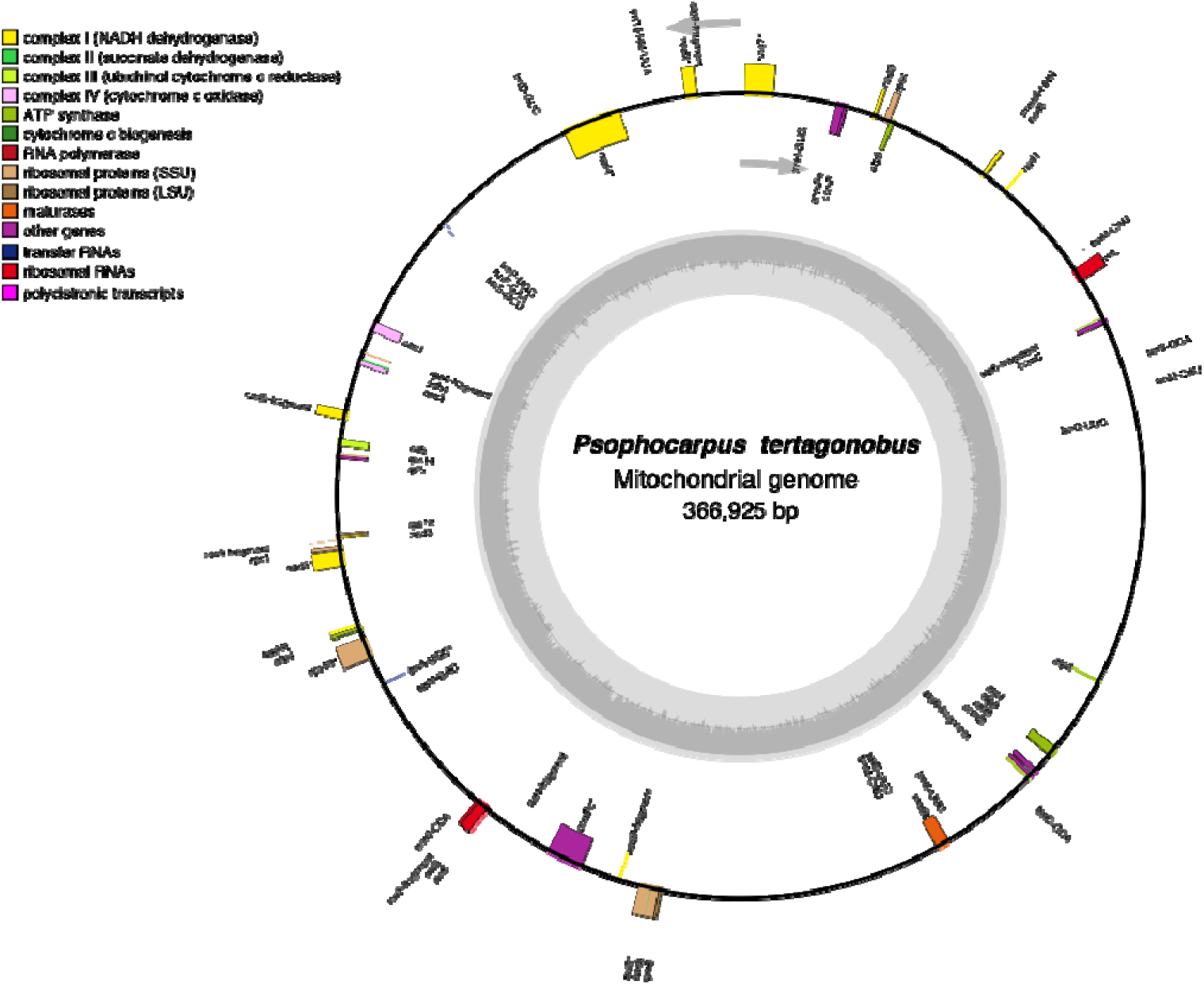
Annotated mitochondrial genome of P. tetragonolobus. The circular diagram represents the complete mitochondrial genome spanning 366,925 bp. Key functional gene categories are color-coded, including polycistronic transcripts, ribosomal RNAs, transfer RNAs, maturases, ribosomal proteins (LSU and SSU), RNA polymerase subunits, and components of respiratory chain complexes I-IV, ATP synthase, and cytochrome c biogenesis. Additional details, such as partial or fragmented genes, are labeled where applicable. The outermost circle displays the genomic arrangement with specified gene loci and their orientations.

The mitochondrial genome contains 64 genes, including 38 protein-coding genes, 12 tRNA genes, and 6 rRNA genes (Fig 1, Table 1). Among the protein-coding genes, key groups include ATP synthase subunits, NADH dehydrogenase subunits, cytochrome genes, and ribosomal protein genes (Fig 1). The mitochondrial genome contains 38 protein-coding genes, which are categorized into distinct functional groups (Table 1). The ATP synthase subunits are represented by 5 genes (atp1, atp4, atp6, atp8, and atp9), encoding components of the ATP synthase complex that drive ATP production through oxidative phosphorylation. The NADH dehydrogenase subunits are encoded by 11 genes (nad1, nad2, nad3, nad4, nad4L, nad5, nad6, nad7, nad9, nad11), which are essential components of Complex I in the electron transport chain, playing a key role in transferring electrons and establishing the proton gradient for ATP synthesis. The cytochrome genes include 4 genes (cox1, cox2, cox3, and cob), which encode subunits of cytochrome c oxidase (Complex IV) and cytochrome b (Complex III), critical for electron transfer and maintaining the mitochondrial energy cycle. Additionally, the genome includes ribosomal protein genes, with 7 genes (rps3, rps4, rps12, rpl2, rpl5, rpl16, and rpl10) encoding structural components of mitochondrial ribosomes required for efficient protein synthesis within mitochondria. Interestingly, 14 fragmented protein-coding genes are presented in the mitogenome, reflecting the dynamic nature of mitochondrial genome organization. These fragmented genes include nad4, nad7, nad2 (with a fragmented form, nad2-fragment), nad5 (with nad5-fragment), nad1 (with nad1-fragment), atp6 (with fragmented forms atp6-1-fragment and atp6-2-fragment), rps4-fragment, rps10, rps3, and ccmFc. The 12 tRNA genes ensure efficient mitochondrial translation, while the 6 rRNA genes, likely reflecting a plant mitochondrial origin, may include fragmented or duplicated forms typical of plant mitochondria (Table 1).

**Table 1:**
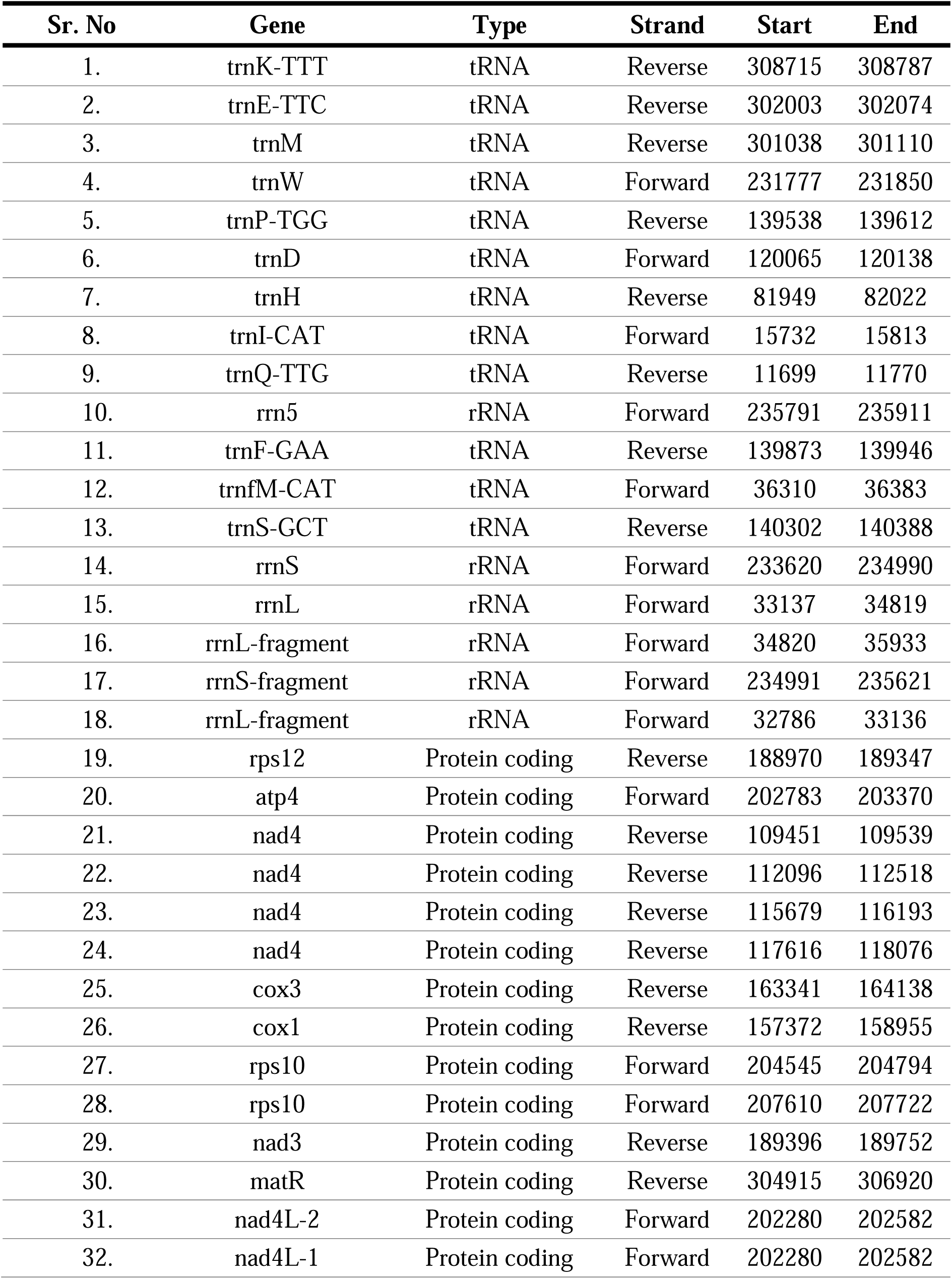
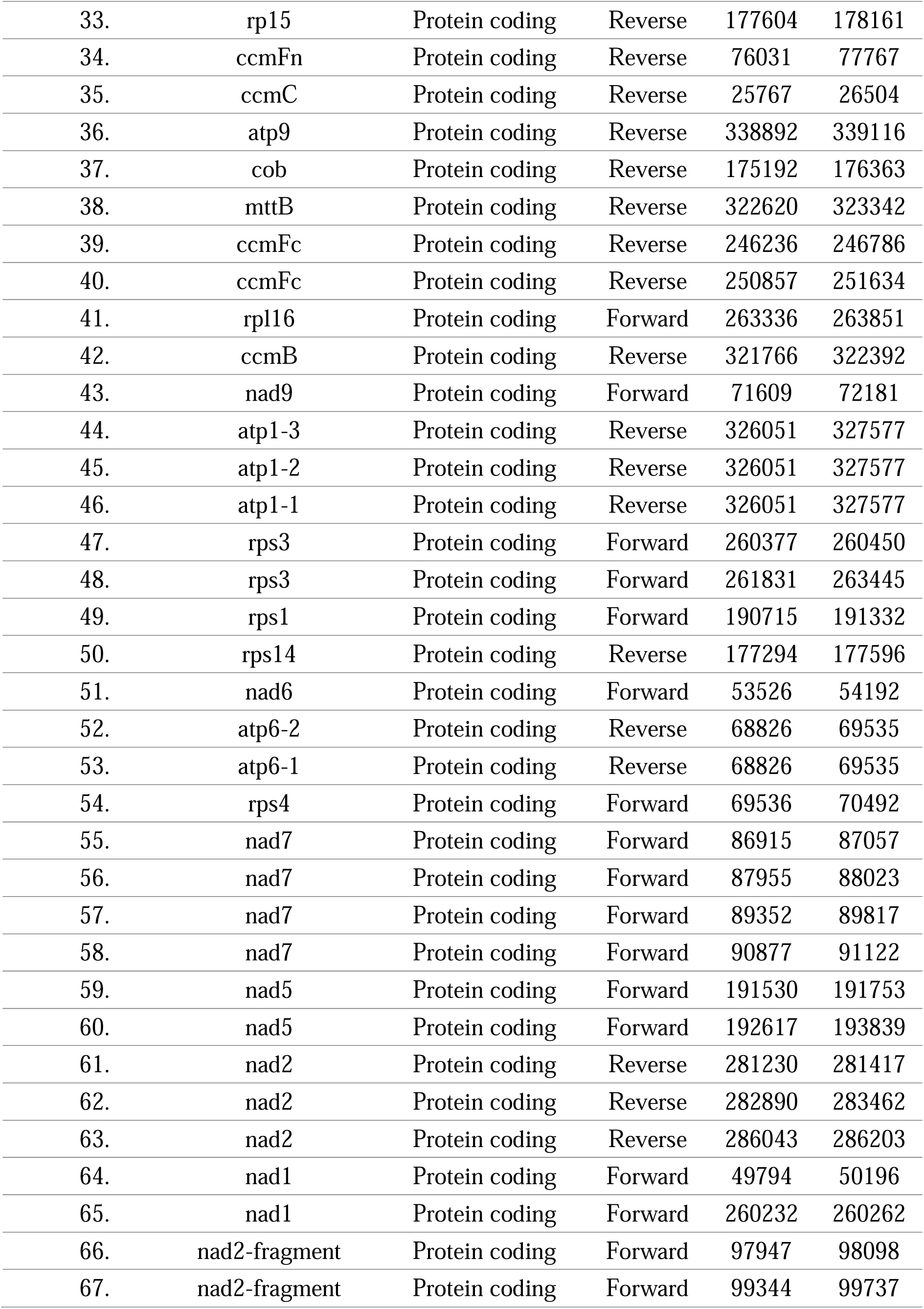
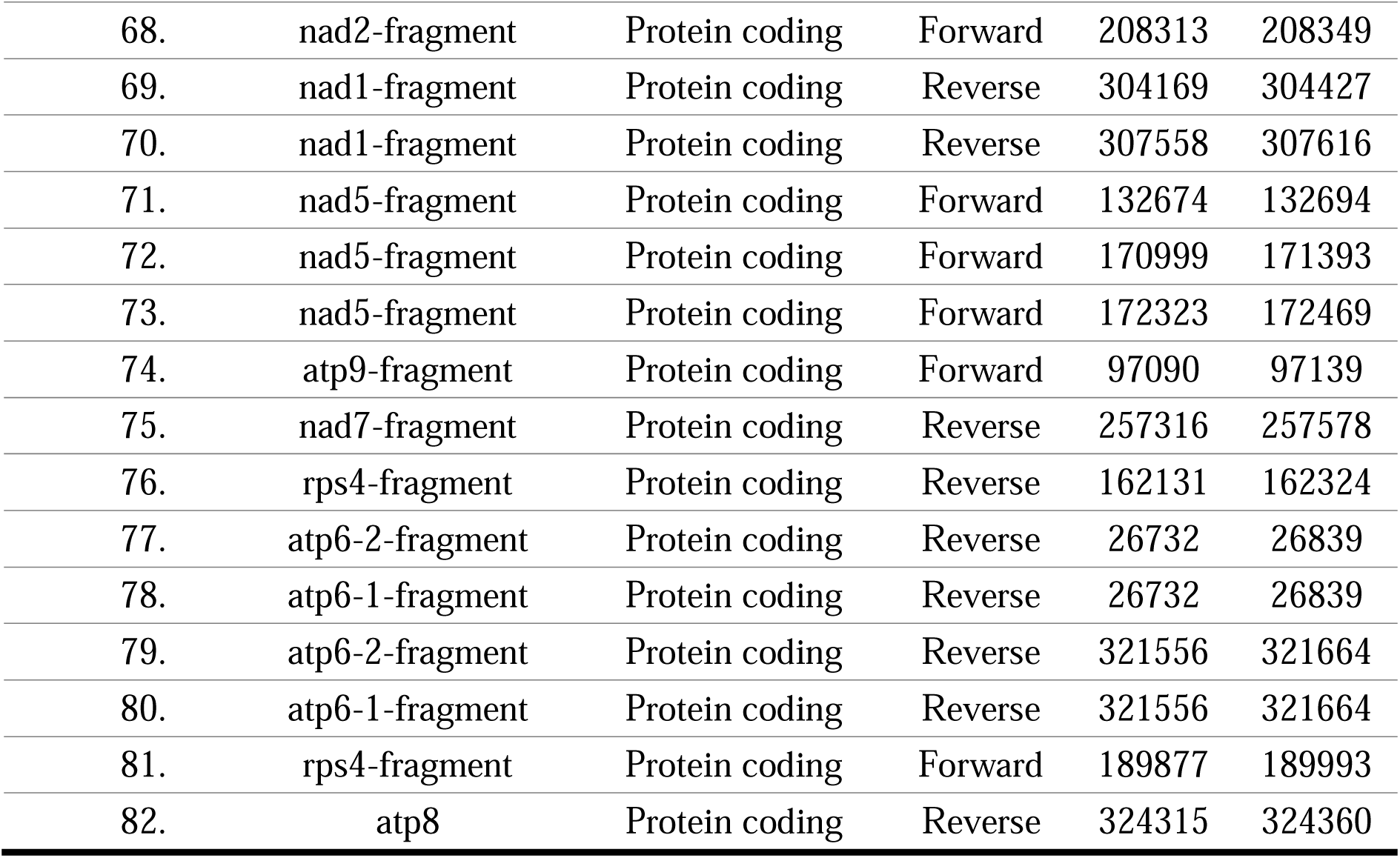
List of Annotated genes in the mitochondrial genome of *P. tetragonolobus*.

### Repetitive Sequences and Large Repeat Regions

The mitochondrial genome of the winged bean contains a significant number of repeated sequences, as illustrated in Fig. 2. The genome includes 100 large repeat sequences spread across the mitogenome, ranging in length from 30 to 110 bp, which play a critical role in structural rearrangements (Fig. 2a-c, Supplementary Table S2). The overall mitochondrial genome has a GC content of 45.44%, while the average GC content of the repeat regions is slightly higher at 47.41% (Supplementary Table S2). These repeats comprise 40 forward repeats and 60 palindromic repeats (Fig. 2b). The longest forward and palindromic repeats both measure 108 bp (Supplementary Table S2). The cumulative length of these dispersed repeats is 18,157 bp, accounting for approximately 4.95% of the total mitochondrial genome (366,925 bp). The analysis revealed that repeats are distributed across both gene-overlapping and intergenic regions in the mitochondrial genome (Fig 2d, Supplementary Table S2). Of the 100 identified repeats, 51 overlap with gene regions, including protein-coding genes such as atp6-1, atp6-2, and nad1, as well as their fragmented forms (Fig 2d, Supplementary Table S2, Supplementary Fig S2). These overlaps highlight the potential functional or structural significance of repeats within coding regions. The remaining 49 repeats are located in intergenic regions (Supplementary Fig S2). The analysis also revealed distinct patterns in the GC content of repeats across the mitochondrial genome. Gene-overlapping repeats (51 total) are predominantly higher in GC content. These repeats align closely with genes, suggesting potential functional or structural roles. In contrast, intergenic repeats (49 total) exhibit broader variation in GC content and are distributed across non-coding regions (Fig 2d).

**Figure 2:**
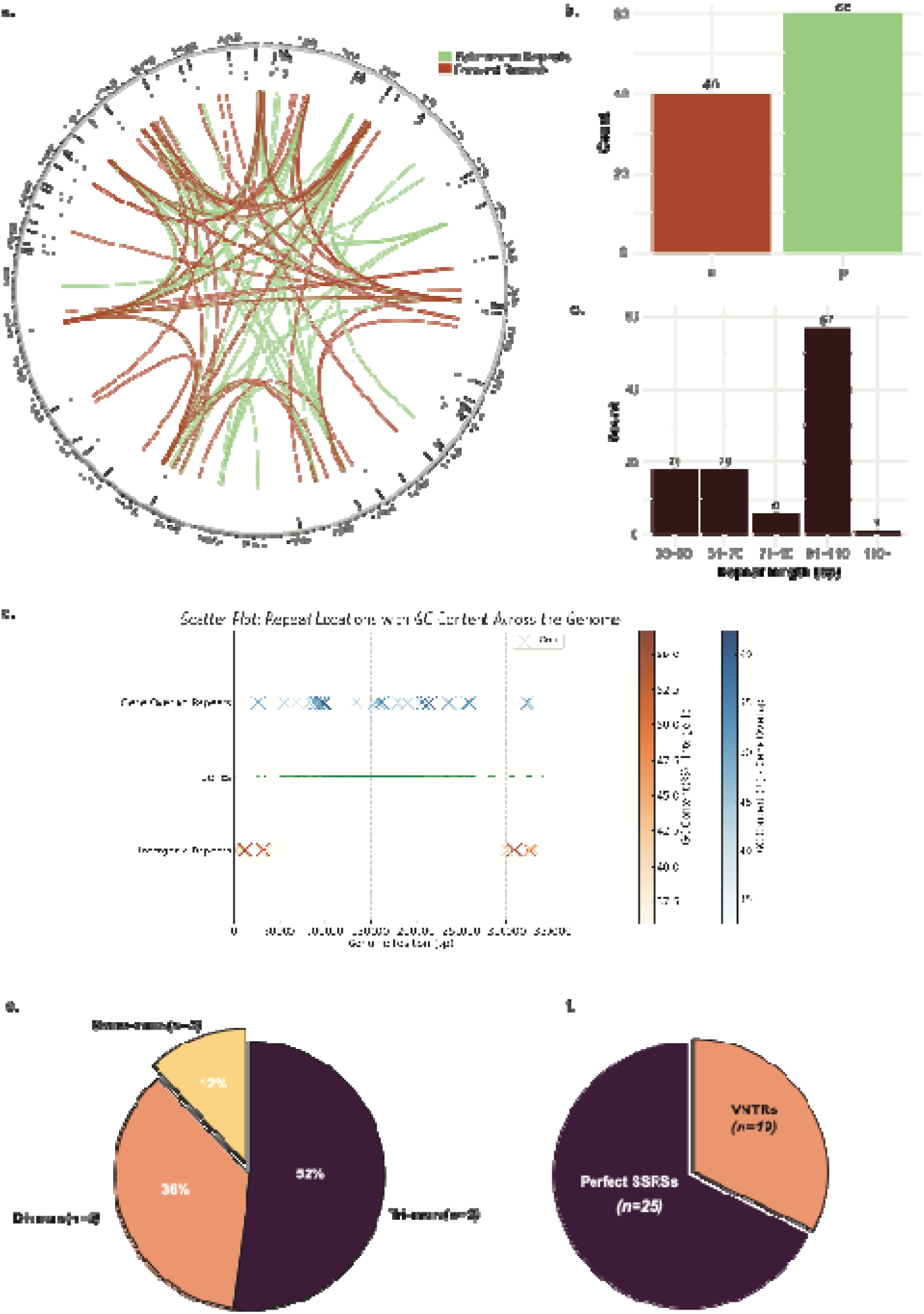
Repeat analysis of the P. tetragonolobus mitochondrial genome. (a) Repeat summary illustrating the distribution of repeat types across the genome, including forward, palindromic, and tandem repeats, categorized by length (bp). (b) Distribution of short sequence repeats (SSRs) and variable number tandem repeats (VNTRs) across genome positions. (c) Percentage breakdown of repeat types in the genome, showing the proportions of perfect SSRs, VNTRs, and other repeat categories. (d-f) Detailed characterization of repeat units by size, including mono-, di-, and tri-mers. Repeat loci and their frequencies are mapped to specific regions of the genome.

**Figure 3:**
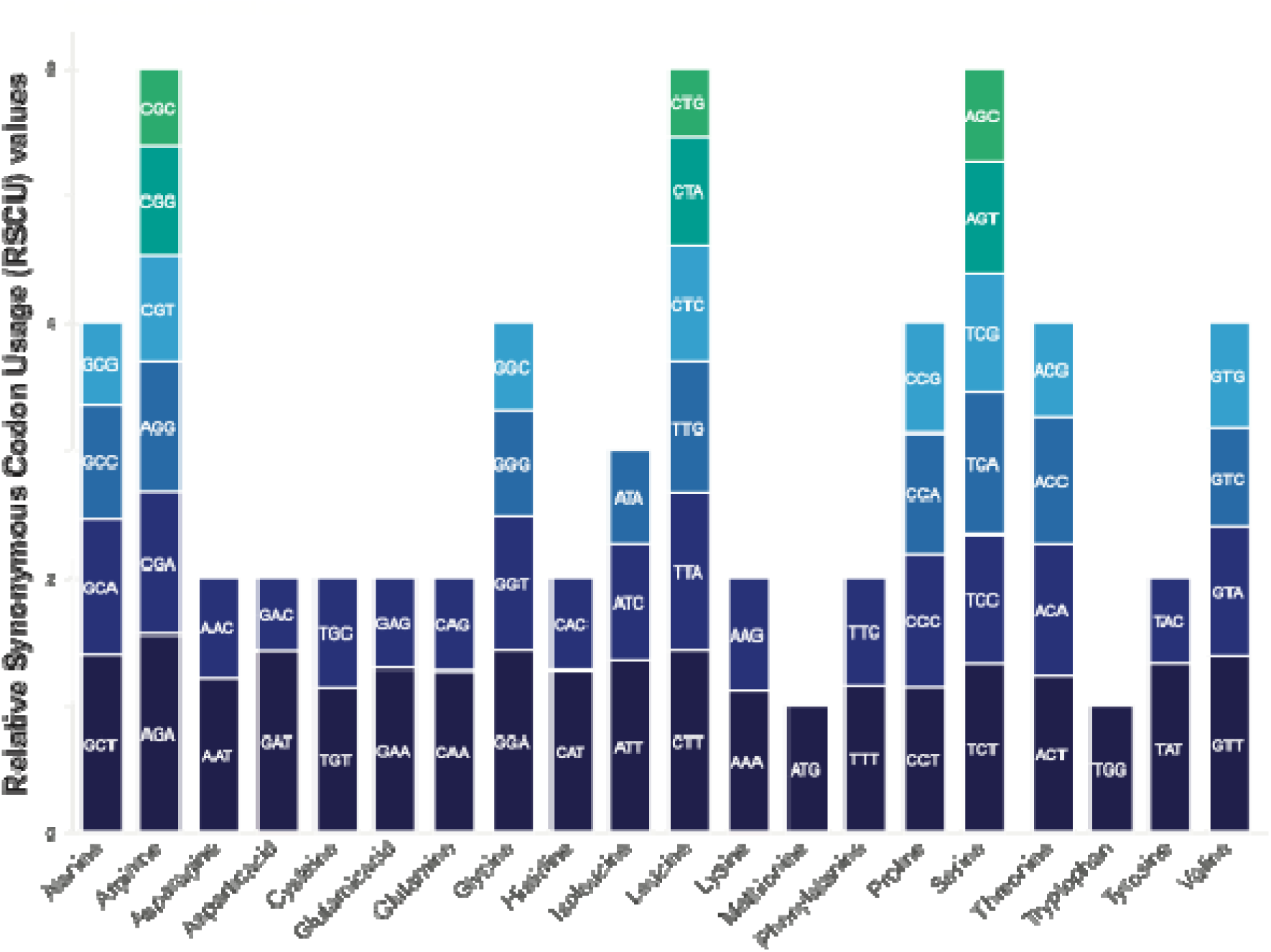
Codon usage and relative synonymous codon usage (RSCU) in the *P. tetragonolobus* mitochondrial genome. The chart illustrates the frequency of each codon across the genome, categorized by amino acids. Codons are grouped by their corresponding amino acids, and their relative synonymous codon usage (RSCU) values are displayed.

A total of 25 simple sequence repeats (SSRs) and 19 variable number tandem repeats (VNTRs) were identified (Fig 2f). Among the SSRs, 3 (12%) are monomers, 9 (36%) di-mers, and 13 (52%) tri-mers, showcasing the diversity of repeat motifs (Fig 2e). Additionally, 19 tandem repeats were identified, with sequence matches of ≥85% and lengths ranging from 10 to 39 bp.

### Codon Usage Bias Analysis

The codon usage analysis of the mitochondrial genome of *P. tetragonolobus* revealed significant codon bias, with distinct preferences for specific codons and amino acids. Among the most frequently used codons were TTT (Phenylalanine) and ATT (Isoleucine), with counts of 392 and 300 and RSCU values of 1.15 and 1.347, respectively, highlighting their preferential usage in mitochondrial protein synthesis. Conversely, codons such as CTG (Leucine) and GAC (Aspartic acid) were underutilized, with low RSCU values of 0.535 and 0.573. Leucine, encoded by six codons, showed a strong bias for CTT (RSCU = 1.429) and TTA (RSCU = 1.234), while CTG was significantly underrepresented. Similarly, serine exhibited a preference for TCT (RSCU = 1.32) and TCA (RSCU = 1.116), indicating selective use of these codons. Stop codons displayed a clear hierarchy of usage, with TGA (RSCU = 1.126) being the most frequent, followed by TAA (RSCU = 1.025) and TAG (RSCU = 0.849). Codons with RSCU values >1, such as GAT (Aspartic acid, RSCU = 1.427) and GAA (Glutamic acid, RSCU = 1.289), were overrepresented, whereas those with values <1, such as GCG (Alanine, RSCU = 0.637), were underutilized, reflecting translational optimization. Overall, the codon usage bias reflects mitochondrial-specific selective pressures, favoring translational efficiency and compatibility with mitochondrial tRNA pools, likely optimizing protein synthesis and reducing translational errors.

### Phylogenetic Analysis

The phylogenetic analysis of the mitochondrial genome of *P. tetragonolobus* reveals its evolutionary relationships within the Fabaceae family and its comparison with tuber crops (Fig 4). Tuber crops were included in the analysis due to the winged bean’s similarity to both legumes and tuber crops. The tree demonstrates clear clustering within the Fabaceae family, with species such as *M. truncatula*, *Trifolium* spp., *L. japonicus*, *P. vulgaris*, *Vigna* spp., and *Glycine* spp. forming well-supported clades, as indicated by bootstrap values ranging from 0.79 to 1.0 (Fig 4). *P. tetragonolobus* clusters closely with legumes such as *G. max* (bootstrap value: 0.97) and *V. radiata* (0.93), underscoring their shared lineage within the Phaseoleae tribe. Its relationship with *G. soja* is similarly well-supported (bootstrap value: 0.86), reflecting genetic similarity among Phaseoleae members. Further divergence is observed within the Fabaceae clade, with *P. tetragonolobus* grouping more distantly with *L. japonicus* (bootstrap value: 0.79) and *T. pratense* (0.73). Its evolutionary distance from *M. truncatula* is marked by a bootstrap value of 0.69, indicating differentiation at the tribal level within Fabaceae. Outside the Fabaceae family, *P. tetragonolobus* exhibits clear divergence from tuber crops, including *S. tuberosum* (bootstrap value: 0.58) and *I. batatas* (0.52). These tuber crops form distinct clusters, highlighting their evolutionary distance from legumes. Similarly, *A. thaliana* and *B. carinata* (bootstrap values: 0.63 and 0.61, respectively) serve as outgroup species, further illustrating the evolutionary separation between Fabaceae and other angiosperms.

**Figure 4:**
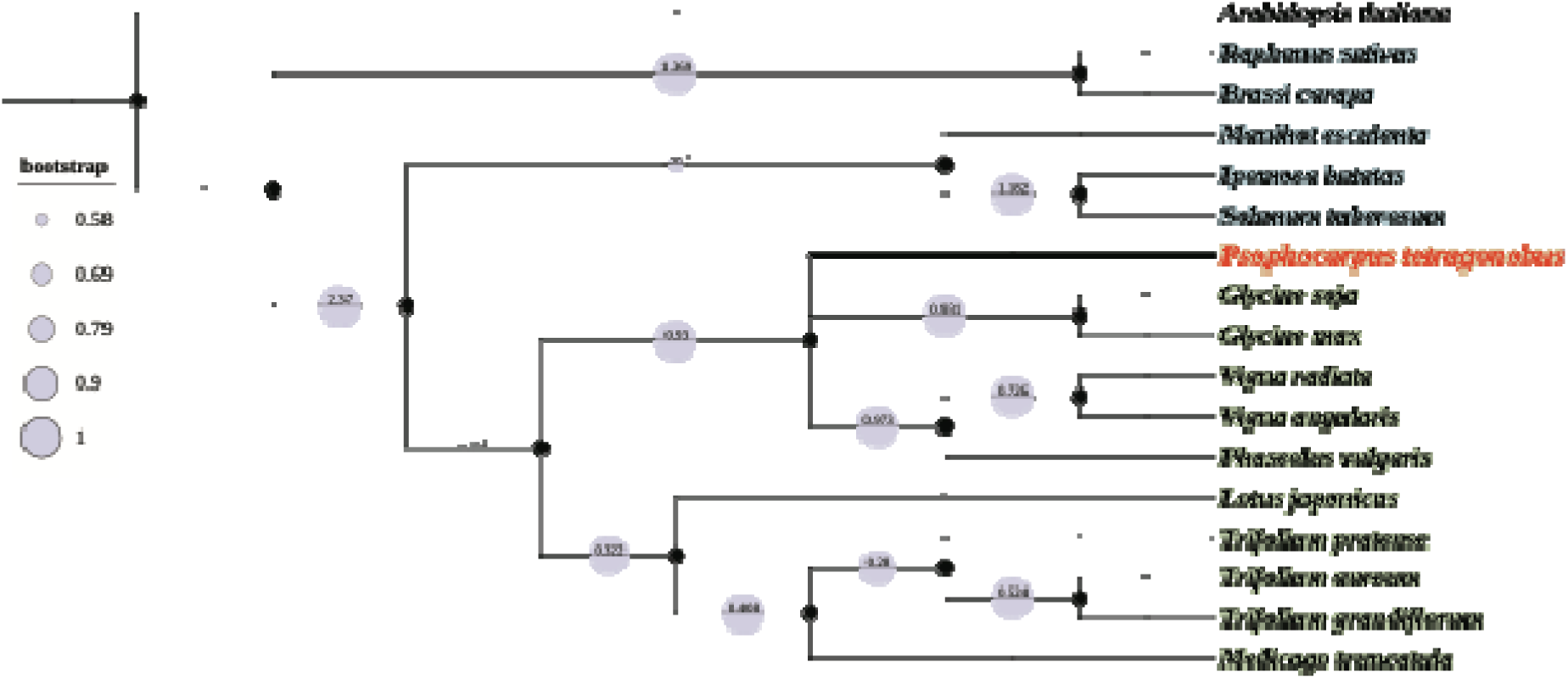
Phylogenetic analysis of *P. tetragonolobus* based on mitochondrial genome sequences. The phylogenetic tree illustrates the evolutionary relationships of *P. tetragonolobus* with other plant species, including legumes (*M. truncatula, Trifolium spp., P. vulgaris, Vigna spp., and Glycine spp.*) and other tuber crops (*S. tuberosum, I. batatas, and M. esculenta*). *A. thaliana* is used as the outgroup to root the tree. Bootstrap values at the nodes indicate the confidence levels for each clade. The analysis confirms the placement of *P. tetragonolobus* within the legume family. The scale bar represents genetic distance.

### Synteny analysis

Synteny analysis using pairwise BLAST results were conducted to evaluate genomic relationships and conservation among *P. tetragonolobus, G. max, V. radiata, P. vulgaris, M. truncatula* (Fig 5). These analyses revealed key conserved and variable regions, providing insights into evolutionary dynamics (Figure 5). The mitochondrial genome of *G. max* (401,300 bp) showed 92% syntenic coverage with *P. tetragonolobus*, characterized by conserved blocks ranging from 10 kb to 150 kb (Supplementary table S2). *V. radiata* (379,425 bp) and *P. vulgaris* (377,980 bp) shared 88% and 86% syntenic coverage, respectively, with conserved blocks spanning 20 kb to 120 kb. Notably, synteny between *P. vulgaris* and *V. radiata* included large, uninterrupted conserved blocks (Fig 5). This level of conservation reflects a shared evolutionary history and strong functional constraints on mitochondrial gene order (Figure 5) and Phylogeny (Figure 4). In contrast, *M. truncatula* (317,280 bp) exhibited fragmented synteny, with only 65% of the *P. tetragonolobus* genome covered by smaller conserved blocks of 5 kb to 50 kb.

**Figure 5:**
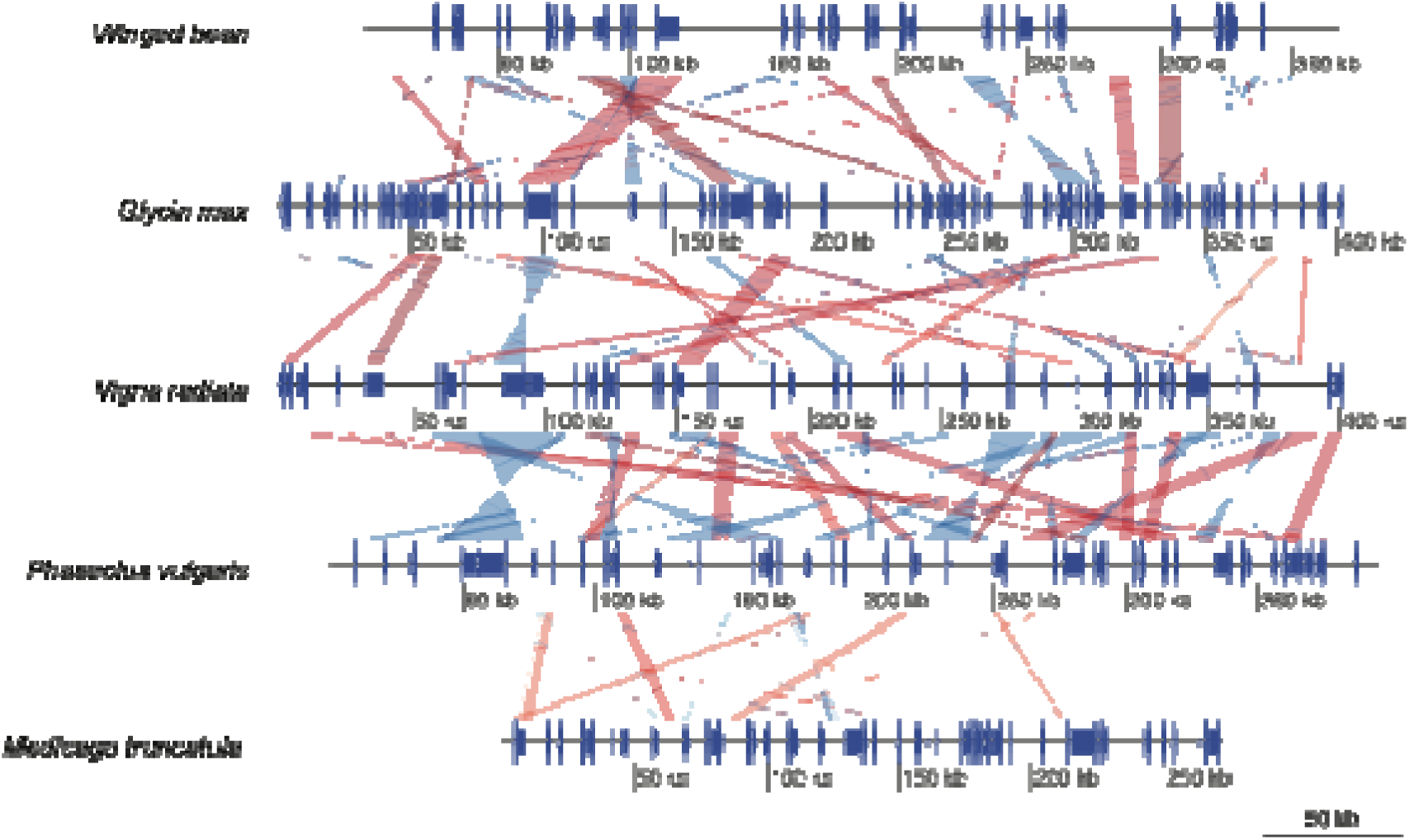
Comparative synteny analysis of the *P. tetragonolobus* mitochondrial genome. The synteny blocks illustrate the conserved genomic regions between the mitochondrial genome of *P. tetragonolobus* (winged bean) and those of other legume species, including *G. max, V. radiata, P. vulgaris*, and *M. truncatula*. Each horizontal bar represents the genomic structure of the respective species, with colour-coded blocks indicating homologous regions.

Percent identity distributions revealed that across all datasets, alignments predominantly fell within the 90% to 98% range, indicating moderate to high conservation. The *P. tetragonolobus vs. G. max* dataset peaked sharply at 98% with also the lowest E-value, while *G. max vs. V. radiata* had a similar peak at 97.8% with E-value below −150 (Supplementary Fig S3).

### Selection pressure analysis

The Ka/Ks ratio analysis of mitochondrial genes in *P. tetragonolobus* revealed varying selective pressures across different genes and species comparisons, providing insights into their evolutionary constraints (Fig 6). For atp1, comparisons with closely related species such as *G. max* and *G. soja* showed extremely low Ka/Ks ratios (0.0249), indicating strong purifying selection, while more distantly related species like *A. thaliana* (0.4100) and *R. sativus* (0.4028) exhibited higher ratios, suggesting relaxed purifying selection (Fig 6). Moderate Ka/Ks ratios were observed with *V. radiata* (0.3007) and *P. vulgaris* (0.3328), reflecting functional constraints in these lineages. The atp4 gene displayed Ka/Ks values close to 1 in comparisons with *B. rapa* (0.9563) and *V. radiata* (0.9093), suggesting neutral or relaxed selection. In contrast, comparisons with *G. max* and *G. soja* showed a Ka/Ks ratio of 0, indicating complete conservation. For atp9, strong purifying selection was predominant, with Ka/Ks ratios well below 1 for most comparisons, though higher values were observed with *L. japonicus* (0.6666) and *T. grandiflorum* (0.3347), reflecting varying levels of divergence. The ccmB gene showed the widest range of Ka/Ks ratios, including evidence of positive selection or relaxed constraints with *M. esculenta* (2.2739) and moderate selection with *A. thaliana* (1.1902) and *R. sativus* (1.3572). In contrast, comparisons with legumes like *G. max* (0.4999) and *V. radiata* (1.1819) suggested stronger purifying selection. Lastly, the ccmC gene exhibited low Ka/Ks ratios across most species, highlighting strong conservation, with exceptions in comparisons with *B. rapa* (0.4466) and *I. batatas* (0.4516), which indicated moderate purifying selection. These findings collectively demonstrate that mitochondrial genes in *P. tetragonolobus* are subject to strong purifying selection, particularly when comparing closely related species. At the same time, more relaxed or adaptive evolutionary pressures are observed in comparisons with distantly related species such as *M. esculenta* and *R. sativus* (Fig 6). The results underscore the evolutionary importance of maintaining mitochondrial gene functionality across lineages.

**Figure 6:**
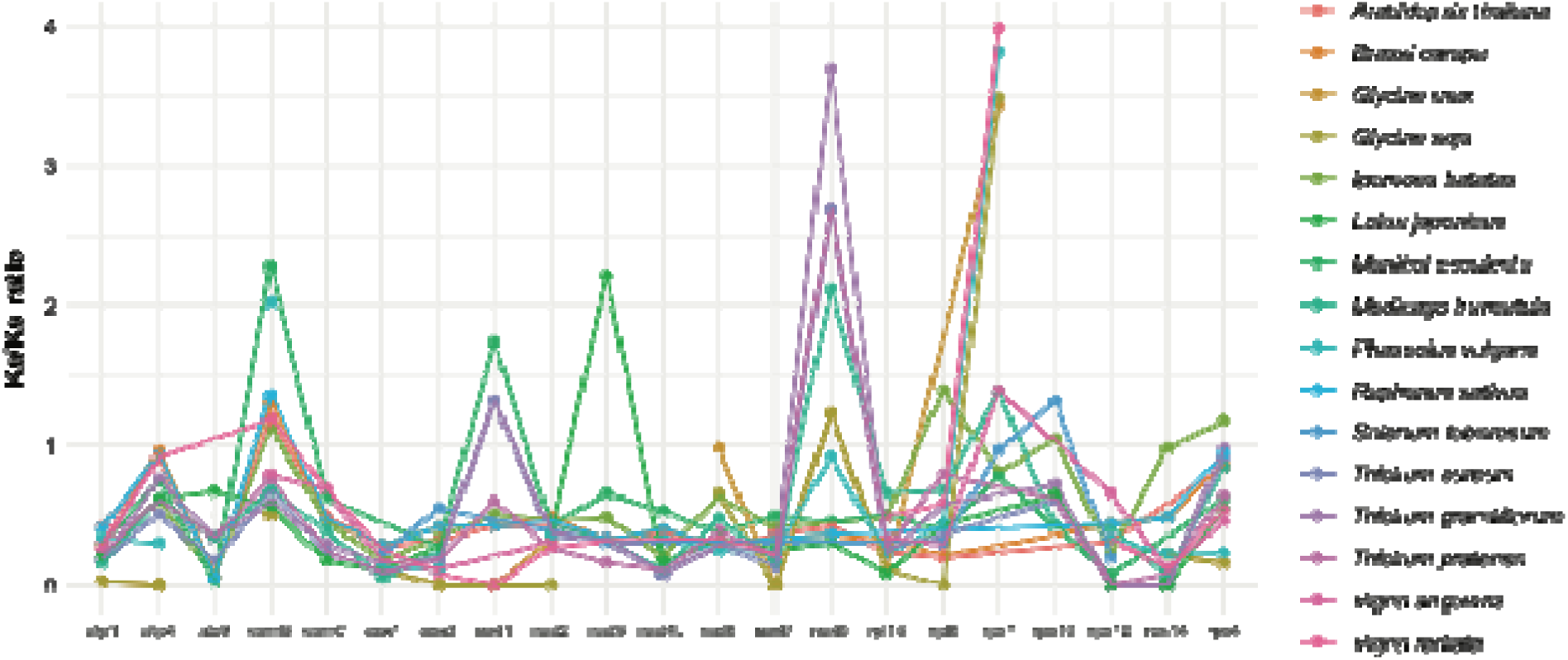
Ka/Ks ratio analysis of protein-coding genes in the *P. tetragonolobus* mitochondrial genome. The chart displays the nonsynonymous to synonymous substitution (Ka/Ks) ratios for protein-coding genes, comparing *P. tetragonolobus* to related plant species, including *A. thaliana, G. max, M. truncatula, V. radiata,* and others. Each bar represents a specific gene’s Ka/Ks ratio, indicating the level of selective pressure acting on it. Ratios below 1 suggest purifying selection, while values approaching or exceeding 1 indicate neutral or positive selection respectively.

### Comparative analysis of winged bean Chloroplast and mitochondrial genomes

The relationship between the effective number of codons (ENC) and GC content at the third codon position (GC3s) was analyzed for mitochondrial and chloroplast genes, as shown in Figure 7a. The ENC values, which range from 20 (extreme codon bias) to 61 (no codon bias), were plotted against GC3s to evaluate codon usage patterns. The expected curve under random codon usage provided a baseline for comparison. The results revealed that the majority of mitochondrial genes clustered closer to the expected ENC curve, with ENC values ranging from 40 to 60 and GC3s spanning 0.25 to 0.75. This distribution indicates moderate codon bias in mitochondrial genes, likely driven by mutational pressures rather than strong selection. In contrast, chloroplast genes exhibited more scattered patterns, with ENC values ranging from 30 to 55 and GC3s between 0.00 and 0.75, suggesting higher variability in codon usage among these genes. The closer alignment of mitochondrial genes to the ENC curve indicates codon usage is predominantly influenced by neutral evolutionary forces, with some contribution from GC content variability. Conversely, the broader spread of chloroplast genes away from the expected curve suggests additional factors, such as translational selection or functional constraints, contribute to their codon usage patterns. These findings highlight distinct codon usage dynamics between mitochondrial and chloroplast genes in *P. tetragonolobus*.

**Figure 7:**
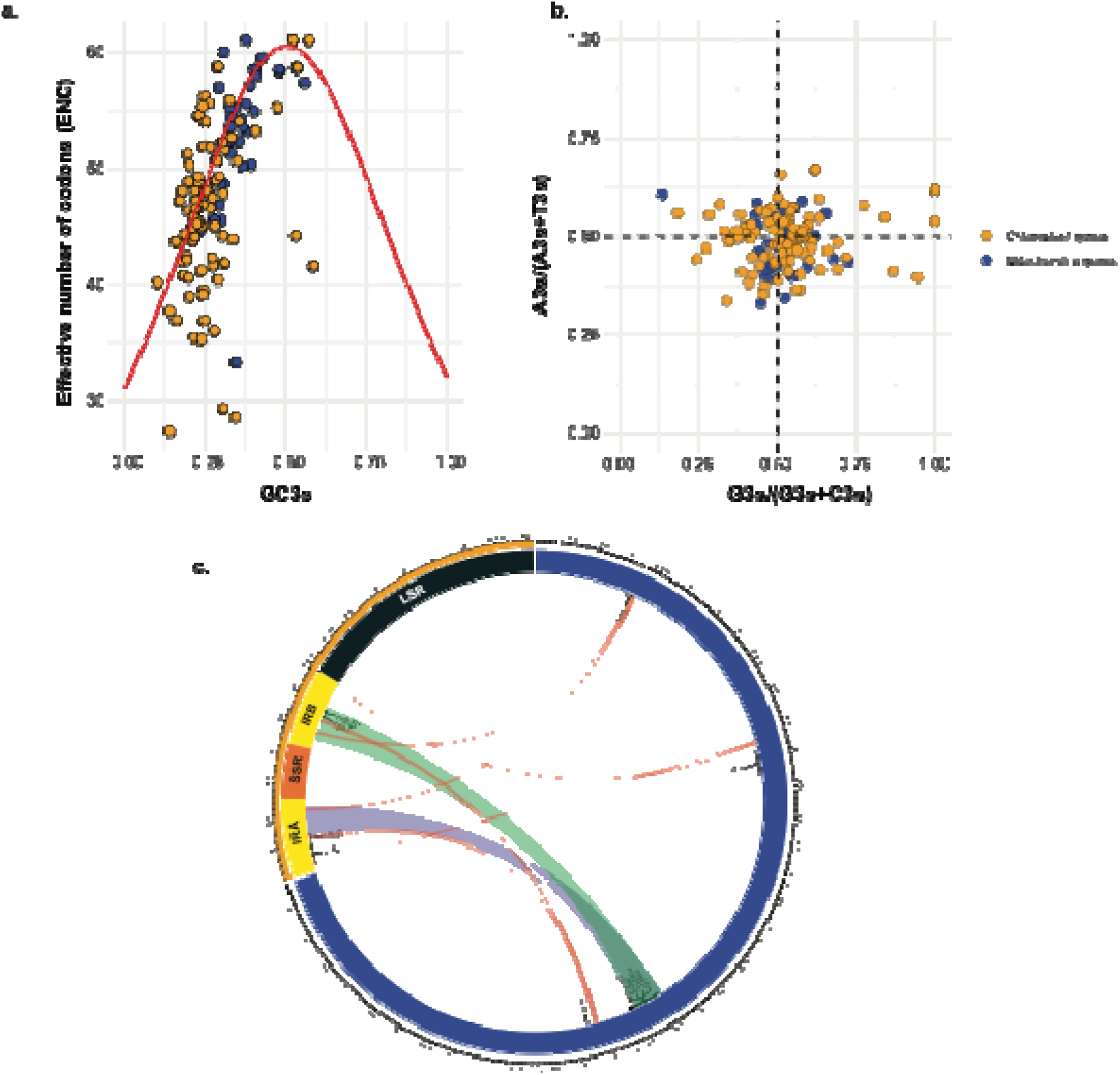
Codon usage bias and genomic features of *P. tetragonolobus* mitochondrial and chloroplast genes. (a) The relationship between GC content at the third codon position (GC3s) and the effective number of codons (ENC) shows a codon usage bias for mitochondrial and chloroplast genes. (b) GC content ratios (G3s/(G3s+C3s) and A3s/(A3s+T3s)) for mitochondrial and chloroplast genes, highlighting compositional biases. (c) Genomic distribution of key genes, including mitochondrial genes (nad4, nad5, rps10) and chloroplast genes (ycf2, ndhB, rps7), along with structural features like inverted repeats (IRA, IRB).

Parity rule 2 (PR2) analysis was conducted to examine the balance between purines (A and G) and pyrimidines (T and C) at the third codon position in the mitochondrial and chloroplast genes of *P. tetragonolobus* (Fig. 7b). The plot displays the ratios A3/(A3 + T3) and G3/(G3 + C3), providing insights into mutational and selective pressures shaping codon usage. The results showed that mitochondrial genes predominantly clustered near the central point (0.5, 0.5), with G3/(G3 + C3) ratios ranging narrowly between 0.4 and 0.6 and A3/(A3 + T3) ratios primarily between 0.45 and 0.55 (Fig 7b). This clustering near equilibrium suggests that mutational pressures dominate over selection in shaping the codon usage of mitochondrial genes. The relatively balanced composition indicates a lack of significant bias in nucleotide usage at the third codon position for these genes. In contrast, chloroplast genes exhibited a much wider distribution across the plot, with G3/(G3 + C3) values ranging from 0.25 to 0.75 and A3/(A3 + T3) spanning from 0.3 to 0.7 (Fig 7b). Many chloroplast genes deviated significantly from the central point, reflecting compositional imbalances at the third codon position. These deviations suggest that both mutational biases and selection pressures play substantial roles in shaping chloroplast gene codon usage. Specifically, the higher variability in chloroplast genes likely results from selective constraints related to their functional roles in photosynthesis and other essential processes.

The circular synteny plot comprehensively compares the mitochondrial and chloroplast genomes of *P. tetragonolobus*, emphasising homologous regions, gene annotations, and inversion synteny (Fig 7c). Key mitochondrial genes located near syntenic regions include trnE-TTC (302,003– 302,074, 94.87% identity, 78 bp alignment), trnM (301,038–301,110, 94.87% identity, 78 bp alignment), trnW (231,777–231,850, 95.91% identity, 98 bp alignment), and trnD (120,065– 120,138, 98.78% identity, 82 bp alignment). In the chloroplast genome, homologous regions align with genes like trnH and trnP, with highly conserved regions such as IR-A and IR-B displaying inversion synteny. These inversions, exemplified by loci such as ndhB and rrnL, showcase rearranged alignments between the genomes, visually represented by reversal in the synteny links (Fig 7c).

Quantitatively, the synteny links reveal high conservation, with percent identities predominantly exceeding 90%. Alignment lengths vary significantly, ranging from smaller functional regions (∼80 bp) to larger conserved regions extending beyond 300 bp (Fig 7c). The visualization incorporates gene names placed adjacent to the genome arcs, alignment statistics, and synteny information, effectively showcasing conserved and inverted genomic architecture. This detailed representation provides valuable insights into the structural and evolutionary relationships between the organellar genomes.

## Discussion

The successful assembly and annotation of the mitochondrial genome of the winged bean mark a significant advancement in the genomic understanding of this underutilized crop. With its total size of 366,925 bp and inclusion of 64 genes, comprising 38 protein-coding genes, 12 tRNA genes, and 6 rRNA genes, the winged bean mitochondrial genome reveals several features characteristic of angiosperm mitochondrial genomes. These findings provide valuable insights into the molecular underpinnings of energy metabolism, stress tolerance, and evolutionary dynamics in this nutritionally and agriculturally important legume.

### Structural Complexity of the Winged Bean Mitochondrial Genome

The variability in mitochondrial genome size among plants reflects a delicate balance between maintaining essential functions and adapting to environmental challenges. Despite the variance in mitogenome sizes, ranging from approximately 66 kb (Skippington et al. 2015) to over 11 Mb (Sloan et al. 2012) across plant species, the functional genes in mitochondrial genomes are highly conserved (T. Kubo & Newton 2008). These conserved genes primarily encode components of the electron transport chain, ATP synthesis, and mitochondrial translation machinery, ensuring the core energy-producing functions of mitochondria remain intact (Adams et al. 2002; Wee et al. 2022). In *P. tetragonolobus*, protein-coding genes are categorized into key groups: ATP synthase subunits, NADH dehydrogenase subunits, cytochrome-related genes, and ribosomal protein genes. These genes encode enzymes and structural proteins vital for oxidative phosphorylation and ATP production, highlighting the functional specialization of mitochondria in energy metabolism and cellular homeostasis.

The sizeable mitochondrial genome sizes observed in plants are often attributed to extensive repetitive sequences, which drive homologous recombination (A. Maréchal & Brisson 2010; Kühn & Gualberto 2012). In *P. tetragonolobus*, 100 repeat sequences, including 40 forward and 60 palindromic repeats, underscore the potential recombination’s role and thus help shape the mitochondrial genome’s structure. Homologous recombination events generate subgenomic molecules and facilitate structural rearrangements, which are both adaptive and functionally significant (A. Maréchal & Brisson 2010). Similar observations in legumes such as *P. vulgaris* and *M. truncatula* suggest conserved mechanisms of mitochondrial genome evolution within the Fabaceae family (Kovar et al. 2018; A. Maréchal & Brisson 2010; Felsted et al. 1977; Chen et al. 2018). Additionally, identifying simple sequence repeats (SSRs) and variable number tandem repeats (VNTRs) in the winged bean’s mitochondrial genome highlights its structural diversity. These repetitive elements enhance genomic plasticity and serve as valuable molecular markers for population genetics and evolutionary studies (Bidyananda et al. 2024; Merritt et al. 2015).

Comparable to other legumes, the winged bean’s mitochondrial genome exhibits unique features indicative of its adaptability to challenging environments. Studies on related species, such as *G. max* and *V. radiata*, have reported similarly dynamic mitochondrial genome structures, reflecting lineage-specific adaptations to environmental pressures (Chang et al. 2013; Manchekar et al. 2006; Alverson et al. 2011; Sloan et al. 2012; Kühn & Gualberto 2012). Notably, the higher repeat density observed in *P. tetragonolobus* suggests a greater potential for recombination-driven genomic plasticity, which may contribute to its exceptional resilience under tropical conditions. This aligns with findings in *L. japonicus*, where repetitive elements were shown to correlate with increased genomic flexibility and enhanced stress adaptation (Shah et al. 2020; Wynn & Christensen 2019). These characteristics position the winged bean as a promising crop for sustainable agriculture in regions facing adverse environmental conditions.

### Codon usage bias and tRNA evolution in the mitogenome of *P. tetragonolobus*

The codon usage analysis of the mitochondrial genome of *P. tetragonolobus* reveals significant preferences for specific codons, correlating with its tRNA availability. This reflects evolutionary optimization for efficient mitochondrial protein synthesis (Jia & Higgs 2008; Iriarte et al. 2021; Parvathy et al. 2022). Among the most frequently used codons are TTT (Phenylalanine) and ATT (Isoleucine), with RSCU values of 1.15 and 1.347, respectively. These biases towards A/T-ending codons are consistent with patterns observed in other plant mitochondrial genomes, such as those of *G. max* and *A. thaliana* (Sinha et al. 2024; Morton & Wright 2007) and have been previously observed in the chloroplast genome of *P. tetragonolobus* (Singh, Singh, Giddhi, Srivast, et al. 2024). Such preferences are linked to mutational biases and selective pressures favouring translational efficiency. Additionally, the presence of tRNA genes such as trnK-TTT, trnE-TTC, and trnM in *P. tetragonolobus* aligns with the observed codon usage patterns. For example, trnK-TTT supports the preferential usage of TTT codons for phenylalanine, enhancing translational accuracy and efficiency. Similarly, underrepresented codons like CTG (Leucine, RSCU = 0.535) and GAC (Aspartic acid, RSCU = 0.573) may reflect limited availability or functionality of the corresponding tRNAs, as seen in other legumes where translational constraints shape codon bias (Mower et al. 2012). In *A. thaliana*, codon usage biases are similarly influenced by tRNA availability, but distinct codon preferences, such as a higher reliance on G/C-ending codons, highlight species-specific adaptations (Wicke et al. 2011). In contrast, *G. max* shows broader codon usage diversity, reflecting its more complex tRNA repertoire (Smith & Keeling 2015).

Stop codon usage in *P. tetragonolobus* shows a clear hierarchy, with TGA (RSCU = 1.126) being the most frequent, followed by TAA (RSCU = 1.025) and TAG (RSCU = 0.849). This pattern aligns with findings in other plants, where stop codons are optimized for termination efficiency and error minimization (Sloan et al. 2012; Sloan et al. 2017). The preference for TGA is particularly notable and may reflect its compatibility with mitochondrial release factors. In summary, the codon usage bias and tRNA gene content of *P. tetragonolobus* highlight the interplay between evolutionary constraints and translational optimization. These results, consistent with other plant mitochondrial genomes trends, provide insights into the evolutionary dynamics shaping mitochondrial functionality in legumes and other plants.

### Inferring evolutionary dynamics of the winged bean mitochondrial genome

The mitochondrial genome of *P. tetragonolobus* provides valuable insights into its evolutionary trajectory within the Fabaceae family, while also offering a first glimpse into its divergence from tuber crops despite morphological similarities. Phylogenetic analysis reveals robust clustering of *P. tetragonolobus* within the Fabaceae family, closely associating it with *G. max* (bootstrap value: 0.97) and *V. radiata* (0.93) in the Phaseoleae tribe. Its relationship with *G. soja* (0.86) underscores shared ancestry within Phaseoleae and highlights genetic diversity between domesticated (*G. max*) and wild (*G. soja*) soybean species. Divergence from more distantly related Fabaceae, such as *M. truncatula* (0.69) and *L. japonicus* (0.79), reflects differentiation at the tribal level, supported by whole-genome and chloroplast genome phylogenies (Ho et al. 2024; Singh, Singh, Giddhi, Srivast, et al. 2024). Despite morphological similarities, *P. tetragonolobus* is evolutionarily distinct from tuber crops like *S. tuberosum* (0.58) and *I. batatas* (0.52), which form separate clusters indicative of independent evolutionary paths.

Phylogenetic and synteny analyses together provide a comprehensive view of evolutionary dynamics within plant species (Zhao et al. 2021; Gao et al. 2020; Krishnamurthy et al. 2022). Our synteny analysis complements phylogenetic findings, revealing high genomic conservation among *P. tetragonolobus* and closely related legumes. The mitochondrial genome of *G. max* shares 92% syntenic coverage with *P. tetragonolobus*, with conserved blocks spanning 10 kb to 150 kb. Similarly, *V. radiata* (88%) and *P. vulgaris* (86%) exhibit substantial synteny, reflecting strong functional constraints and shared evolutionary history. In contrast, *M. truncatula* displays fragmented synteny, with 65% coverage and smaller conserved blocks (5 kb to 50 kb), signifying more significant divergence.

The Ka/Ks ratio analysis of mitochondrial genes provides critical functional insights, reinforcing patterns observed in synteny and phylogeny. Genes like *atp1* exhibit consistently low Ka/Ks ratios (0.0249) in comparisons with closely related species such as *G. max* and *G. soja*, reflecting strong purifying selection that supports phylogenetic clustering and syntenic conservation in our study. Higher Ka/Ks ratios observed with distantly related species, such as *A. thaliana* (0.4100) and *M. esculenta* (2.2739), suggest relaxed purifying selection or adaptive divergence, consistent with fragmented synteny in these lineages. Similarly, moderate Ka/Ks ratios in comparisons with *V. radiata* (0.3007) and *P. vulgaris* (0.3328) indicate functional constraints in line with their shared evolutionary history. Positive selection signals, such as those observed in *ccmB* with *M. esculenta*, align with functional divergence and phylogenetic distance, whereas conserved genes like *ccmC* with low Ka/Ks ratios emphasize strong evolutionary constraints that maintain essential mitochondrial functions. These findings align with previous studies using organelle genomes in species like Mangifera (Niu et al. 2022), Brassica (Chang et al. 2011), *Punica granatum* L (Lu et al. 2023), *Cucumis sativus (Park et al. 2021)* and other legumes (Singh, Singh, Giddhi, Srivast, et al. 2024; Alverson et al. 2011; Choi et al. 2021; Lee et al. 2021; Choi et al. 2022; Satrio et al. 2023; Yang et al. 2023)

This variation could point to evolutionary processes like domestication and natural selection shaping these lineages (Diamond 2002; Alam & Purugganan 2024). *P. tetragonolobus* diverged into the Phaseoleae tribe alongside *G. max* (soybean) and *V. radiata* (mung bean) (Singh, Singh, Giddhi, Srivast, et al. 2024; Ho et al. 2024) adapting to specific environments with distinct traits—climbing habits in winged beans versus bush growth in soybeans. A similar divergence is observed in grasses like *Oryza sativa* (rice) and *Zea mays* (maize), where rice adapted to water-logged tropical conditions and maize evolved for temperate dryland farming (Birch et al. 2014; Ilic et al. 2003; Chen et al. 2021). These findings collectively emphasize the evolutionary importance of maintaining mitochondrial gene functionality across lineages. The combined phylogenetic, synteny, and Ka/Ks analyses contribute to understanding mitochondrial genome evolution and its role in shaping lineage-specific adaptations. Future studies should emphasize functional analyses to explore the adaptive significance of conserved and divergent genomic regions.

### Evolutionary insights from codon usage and genomic architecture of organelle genomes

The codon usage and synteny analyses of mitochondrial and chloroplast genomes in *P. tetragonolobus* reveal complementary evolutionary dynamics driven by distinct pressures and functional roles. ENC-GC3s analysis shows mitochondrial genes clustering near the expected ENC curve, with moderate codon bias (ENC 40–60, GC3s 0.25–0.75) predominantly shaped by mutational pressures (Wang et al. 2023; Sun et al. 2024; Indrabalan et al. 2021). In contrast, chloroplast genes display greater variability (ENC 30–55, GC3s 0.00–0.75), suggesting additional influences like translational selection and functional constraints related to photosynthesis (Xiao et al. 2024; Rao et al. 2024; Li et al. 2023; Wang et al. 2018). PR2 analysis supports these findings: mitochondrial genes cluster near equilibrium (G3/(G3 + C3) and A3/(A3 + T3) around 0.5), reflecting mutational dominance. Chloroplast genes exhibit broader distributions (G3/(G3 + C3) 0.25–0.75, A3/(A3 + T3) 0.3–0.7), indicating a balance of mutational biases and selective pressures (Paul et al. 2018; Qiu et al. 2011; Morton & Wright 2007; Kimura 1981). In contrast, *A. thaliana* displays a broader range of GC3s values and stronger codon bias for certain genes, reflecting more pronounced selective pressures and functional constraints in its mitochondrial genome (Sloan et al. 2017)

The circular synteny plot highlights high conservation in mitochondrial genes such as *trnE-TTC* and *trnM* (>94% identity), aligning with their balanced codon usage. Chloroplast regions like IR-A and IR-B show inversion synteny and structural rearrangements, reflecting their adaptive variability. Percent identities above 90% across homologous regions demonstrate shared evolutionary origins, while alignment lengths (∼80 to 300 bp) underscore lineage-specific adaptations. Mitochondrial genomes exhibit stability driven by mutational pressures and functional constraints, supporting energy metabolism. Chloroplast genomes show more significant variability, shaped by selection pressures linked to photosynthesis and environmental adaptation. The interplay between codon usage and structural conservation underscores the evolutionary strategies balancing conservation and adaptability in organellar genomes. Future studies should integrate functional data to explore these patterns further (Niu et al. 2022; Allen 2015; Wang et al. 2007; Cui et al. 2021; Zhang et al. 2012; Li et al. 2024).

The mitochondrial genome of *P. tetragonolobus* offers critical insights into the evolutionary dynamics and functional adaptations of organellar genomes. Codon usage patterns highlight the role of mutational pressures in shaping mitochondrial genes, while chloroplast genes display greater variability driven by translational selection and functional constraints. Synteny analysis underscores high genomic conservation, interspersed with lineage-specific structural rearrangements that reflect evolutionary divergence. These findings collectively emphasize the interplay of conservation and adaptability in organellar genomes, shedding light on the molecular mechanisms underlying stress tolerance, energy metabolism, and environmental resilience in this underutilized crop. Future research integrating functional, transcriptomic, and proteomic data will further elucidate the adaptive significance of these evolutionary patterns and support the sustainable utilization of *P. tetragonolobus* in agriculture.

## Materials and Methods

### Sampling and DNA extraction

For the accusation of the mitochondrial genome of winged bean, young leaves were harvested from dual-purpose cultivar AKWB-1 grown in the experimental farm at ICAR-Indian Institute of Agricultural Biotechnology, located in Ranchi, Jharkhand, India, with the geographical coordinates 23°16’27.6“N, 85°20’29.4“E. Before harvesting, AKWB-1 plants were self-pollinated for three consecutive generations to ensure homozygosity. High-quality total genomic DNA was isolated using the cetyltrimethylammonium bromide (CTAB) extraction method. (Doyle 1991). To remove RNA contamination 2.0 μl of RNase A (10 mg/ml, HiMedia) was added to 20 μl of DNA dissolved in TE buffer (Tris–EDTA, pH = 8.0), followed by incubation at 37 °C for 3–4 h. The whole-genome sequencing library was prepared using the TruSeq DNA PCR-Free Library Prep Kit, following the manufacturer’s protocol. After evaluating the library’s quality and quantity, whole-genome sequencing was carried out on an Illumina HiSeq2500 platform using a paired-end (2 × 150 bp) sequencing strategy with the TruSeq Rapid SBS Kit. Simultaneously, the same biomass was used for long-read sequencing with PacBio technology. The library for PacBio sequencing was constructed using the SMRTbell Express Template Preparation Kit 2.0 and purified with AMPure PB beads according to the manufacturer’s instructions. Approximately 80 pM of the library was loaded onto a single SMRTcell with 8M ZMW and sequenced on the PacBio Sequel II system in CCS/HiFi mode.

### Sequencing quality assessment

Illumina short reads were checked for quality using FastQC v. 0.12.0 (Andrews n.d.) and read counts were extracted. Sequencing reads were then trimmed for adapter sequences and sequencing quality using Trimmomatic v. 0.39 (Bolger et al. 2014) using the following settings: illuminaclip=TruSeq3-PE.fa:2:30:10, leading=10, trailing = 10, sliding-window = 5:10 and minlen = 50. The raw subreads obtained from the PacBio Sequel II sequencer were processed to generate highly accurate circular consensus sequences (HiFi reads) using the CCS (Circular Consensus Sequencing) tool (Wenger et al. 2019).

### Mitochondria genome assembly and annotation

We assembled the mitogenome using trimmed and filtered PacBio long reads with the MitoHiFi tool v.3.2.2 (Uliano-Silva et al. 2023). We choose *G. max* (PRJNA927338) as the closest reference as it was shown to be the closest species when comparing the chloroplast genome (Singh, Singh, Giddhi, Srivastava, et al. 2024). The model organism was set to plant with “*-a plant*” option. The most extended assembly with the highest coverage was considered the best mitochondrial genome candidate. We then employed GetOrganelle tool v.1.7.7.1 (Jin et al. 2020) with short reads specifying the embryophyta plastid database and using the candidate mito genome fasta file. The parameters used were “*-k 21,45,65,85,105 -P 1000000 -w 85*”. The Maximum extension rounds were set to 20. The resultant assembly in the graphical fragment assembly (GFA) was manually checked in Bandage (Wick et al. 2015). Contigs with coverage of less than 50 or those that mapped to the chloroplast genome were removed, and the retained graph was exported in fasta format.

Annotation of the mitochondrial genome was then conducted, referencing all previously published mitochondrial genomes of Fabaceae family, utilising Geseq software for this purpose (Tillich et al. 2017). The output genebank and gff files were manually checked, corrected, and a circular representation was drawn using OGDRAW (Greiner et al. 2019). The complete mitochondrial genome sequence of the winged bean was submitted to GenBank with the accession number PQ877725.1.

### Investigation of DNA repeat sequences

For the identification of repetitive sequences within the genome, microsatellites, tandem repeats, and dispersed repeats were conducted using Krait v1.3.3 software (Du et al. 2018), Tandem repeats were analyzed using Tandem Repeats Finder (Benson 1999) and REPuter (Kurtz et al. 2001). Long repeats in the chloroplast genome of the winged bean were specifically identified using REPuter (Kurtz et al. 2001). with repeats categorized as forward (F) or palindromic (P) at a Hamming Distance of 3 and a minimum repeat length of 30. Additionally, the chloroplast genome was examined for perfect SSRs (pSSRs), compound SSRs (cSSRs), and variable number tandem repeats (VNTRs) using the Krait v1.3.3 software (Du et al. 2018). The analysis followed specific criteria: mono-nucleotide repeat motifs required at least 10 repeats, di-nucleotide motifs required a minimum of five repeats, and tri-nucleotide motifs also needed at least five repeats, while tetra-, penta-, and hexa-nucleotide motifs required a minimum of four repeats. If the distance between two SSRs was less than 10 bp, they were classified as cSSRs.

#### Phylogeny and synteny analysis

The nucleotide sequences of all predicted mitochondrial genes of *P. tetragonolobus* and the reported mitochondrial genes for nine other Fabaceae species, including *Vigna radiata, Vigna angularis, Lotus japonicus, Phaseolus vulgaris, Glycine max, Glycine soja, Medicago truncatula, Trifolium pratense, Trifolium aureum, Trifolium grandiflorum,* along with five tuber crop species including *Raphanus sativus, Brassi carapa, Manihot esculenta, Ipomoea batatas, Solanum tuberosum* and *Arabidopsis thaliana* as the outgroup, were obtained from NCBI. Alignment and subsequent phylogenetic analysis were conducted using the genes shared across the species. The sequences were aligned using MAFFT v7.487 (Katoh et al. 2002; Nakamura et al. 2018) with 1000 iterative refinement steps using “--maxiterate 1000”. The resulting aligned sequences were saved in FASTA format. Phylogenetic trees for each aligned gene were inferred using RAxML-NG v.1.2.2 (Kozlov et al. 2019) with 1000 Bootstrap replicates and the “General Time Reversible (GTR) with gamma-distributed rate variation among sites (GTR + G)” model. The best gene tree files were used to prepare a multigene-based species tree with ASTRAL v5.7.8 63 (Mirarab et al. 2014; Zhang et al. 2018). Finally, the phylogenetic tree was visualised using FigTree v1.4.4 (http://tree.bio.ed.ac.uk/software/figtree/).

We then used the mitogenomes of *G. max*, *V. radiata, P. vulgaris, M. truncatula*, and analyzed the synteny and rearrangements in the mitogenomes of *P. tetragonolobus.* For synteny plots, pairwise BLASTN results were generated using the whole mitogenomes of each species in R using a custom script. Information on BLAST hits was visualised in R using the genoPlotR package (Guy et al. 2010). Information on BLAST hits among homologous segments was also visualised in R using the genoplotR package (Guy et al., 2010). BLAST results were also summarised in R.

#### Estimation of adaptive evolution of protein-coding genes

To analyze the degree of non-synonymous (Ka) and synonymous (Ks) substitutions, as well as their ratio (dN/dS), the coding DNA sequences (CDS) of *P. tetragonolobus* were compared with 16 other species including *G. max*, *G. soja*, *V. radiata*, *P. vulgaris*, *L. japonicus*, *M. truncatula*, *T. aureum*, *T. grandiflorum*, *T. pratense*, *B. rapa*, *M. esculenta*, *I. batatas*, *R. sativus*, *S. tuberosum*., along with *A. thaliana*. For this, we performed pairwise alignments using MAFFT v7 (Katoh & Standley, 2013), and then Ka, Ks substitutions and their ratio were calculated using KaKs_Calculator 3 (Zhang 2022) with the MA model.

#### Codon usage analysis

The codon usage within the coding part of the chloroplast was calculated using CodonW v1.4.4 available through https://codonw.sourceforge.net, using universal codon standards. The analysis included T3, C3, A3, and G3, which represent the nucleotide usage frequencies at the third position of codons. GC3s denote the GC content at the third position of synonymous codons, while the effective number of codons (ENc) was calculated using the CodonW software. Additionally, the GC content at the first (GC1), second (GC2), and third (GC3) codon positions was determined using the online CUSP tool available at https://www.bioinformatics.nl/cgi-bin/emboss/cusp.

The Parity Rule 2 (PR2) plot was employed to analyze the nucleotide composition at the third position of codons. This widely used approach assesses whether mutation pressure or selection pressure predominantly influences the nucleotide composition in DNA double strands. In the PR2 plot, the horizontal axis represents G3/(G3+C3), while the vertical axis corresponds to A3/(A3+T3). If mutation pressure alone shapes codon composition, all points will cluster at the plot’s center. However, deviations from the center indicate the influence of natural selection and other factors on codon usage.

In the ENc-GC3s plot, GC3s values are displayed on the horizontal axis, while the actual ENc (actENc) values are plotted on the vertical axis. Codons for methionine (Met) and tryptophan (Trp) were excluded from the analysis because they lack synonymous codons. The expected ENc (expENc) values were calculated using the following formula:

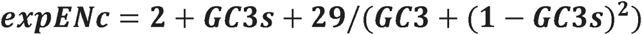

A standard curve was generated based on the expected values. Data points on the standard curve indicate a significant influence of mutational pressure on codon usage. Conversely, data points significantly below the standard curve indicate that natural selection and other factors may be the primary driving forces influencing codon usage patterns (Shi et al., 2024). ENC ratio (ENc ratio) was determined using the following formula, which illustrates the difference between the actENc and expENc values:

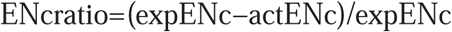

Neutrality plot analysis was performed to assess the relative impact of mutation pressure and natural selection on codon usage patterns. This analysis involved examining GC12 (the average GC content at the first and second codon positions), GC1, GC2, and GC3. In the plot, GC3 values were placed on the horizontal axis, while GC12 values were represented on the vertical axis, creating two-dimensional scatter plots for individual genes. If the regression curve’s slope is near 0 and there is no significant correlation between GC12 and GC3, it suggests that natural selection primarily drives codon usage patterns. On the other hand, if the slope approaches or equals 1 with a strong correlation, it indicates that mutation pressure plays a significant role in shaping gene evolution.

#### Chloroplast to mitochondrion DNA transformation

DNA migration is common in plants and varies from species to species. This phenomenon occurs during autophagy, gametogenesis, and fertilization. The previously published chloroplast genome of the winged bean (Singh, Singh, Giddhi, Srivastava, et al. 2024) was compared with the mitochondrial sequence using Blastn software (Camacho et al. 2009) and visualised using Circoletto (Darzentas 2010) to identify the synteny between the organelle genomes Screening criteria were set as the matching rate ≥ 70%, E-value ≤ 1e ^−10^, and length ≥ 40.

## Data availability

The complete mitochondrial genome sequence of *Psophocarpus tetragonolobus* L. was submitted to GenBank with the accession number PQ877725.1.

## Acknowledgement

We acknowledge the Nucleome Informatics, Pvt. Limited Hyderabad for Illumina and PacBio sequencing.

## Authors Contribution

NKS-data analysis and writing manuscript, BKS-sequencing, and writing the manuscript, PK-data analysis, AP-data analysis, SK-Sample generation and sequencing, SKB-data analysis, APt-Coordinated the project, manuscript editing, VBP-Coordinated the project, manuscript editing, SR-Coordinated the project, manuscript editing, KUT-Planning the experiment, sequencing, and writing the manuscript

## References

Adams, K.L. et al., 2002. Punctuated evolution of mitochondrial gene content: high and variable rates of mitochondrial gene loss and transfer to the nucleus during angiosperm evolution. Proceedings of the National Academy of Sciences of the United States of America, 99(15), pp.9905–9912. Available at: 10.1073/pnas.042694899.

Adams, K.L. & Palmer, J.D., 2003. Evolution of mitochondrial gene content: gene loss and transfer to the nucleus. Molecular phylogenetics and evolution, 29(3), pp.380–395. Available at: 10.1016/s1055-7903(03)00194-5.

Alam, O. & Purugganan, M.D., 2024. Domestication and the evolution of crops: variable syndromes, complex genetic architectures, and ecological entanglements. The plant cell, 36(5), pp.1227–1241. Available at: https://academic.oup.com/plcell/article/36/5/1227/7577726.

Allen, J.F., 2015. Why chloroplasts and mitochondria retain their own genomes and genetic systems: Colocation for redox regulation of gene expression. Proceedings of the National Academy of Sciences of the United States of America, 112(33), pp.10231–10238. Available at: https://www.pnas.org/doi/10.1073/pnas.1500012112#:∼:text=Abstract,events%20in%20photosynthesis%20and%20respiration.

Alverson, A.J. et al., 2011. The mitochondrial genome of the legume Vigna radiata and the analysis of recombination across short mitochondrial repeats. PloS one, 6(1), p.e16404. Available at: https://pmc.ncbi.nlm.nih.gov/articles/PMC3024419/.

Andrews, S., FastQC: A quality control analysis tool for high throughput sequencing data, Github. Available at: https://github.com/s-andrews/FastQC [Accessed January 16, 2025].

Arrieta-Montiel, M.P. & Mackenzie, S.A., 2011. Plant Mitochondrial Genomes and Recombination. Plant Mitochondria, pp.65–82. Available at: https://link.springer.com/chapter/10.1007/978-0-387-89781-3_3 [Accessed December 24, 2024].

Benson, G., 1999. Tandem repeats finder: a program to analyze DNA sequences. Nucleic acids research, 27(2), pp.573–580. Available at: 10.1093/nar/27.2.573.

Bepary, R.H. et al., 2023. Biochemical composition, bioactivity, processing, and food applications of winged bean (*Psophocarpus tetragonolobus*): A review. Legume science, 5(3). Available at: https://onlinelibrary.wiley.com/doi/abs/10.1002/leg3.187.

Bidyananda, N. et al., 2024. Plant genetic diversity studies: Insights from DNA marker analyses. International Journal of Plant Biology, 15(3), pp.607–640. Available at: https://www.mdpi.com/2037-0164/15/3/46.

Birch, J.L. et al., 2014. Phylogenetic investigation and divergence dating of*Poa*(Poaceae, tribe Poeae) in the Australasian region: Australasian*Poa*Phylogenetics and Dating. Botanical journal of the Linnean Society. Linnean Society of London, 175(4), pp.523–552. Available at: https://academic.oup.com/botlinnean/article-abstract/175/4/523/2416406?redirectedFrom=fulltext.

Bock, R. & Knoop, V., 2012. Genomics of Chloroplasts and Mitochondria, Springer Science & Business Media. Available at: https://play.google.com/store/books/details?id=za8Nbr7L87kC.

Bolger, A.M., Lohse, M. & Usadel, B., 2014. Trimmomatic: a flexible trimmer for Illumina sequence data. Bioinformatics, 30(15), pp.2114–2120. Available at: http://www.pubmedcentral.nih.gov/articlerender.fcgi?artid=PMC4103590.

Camacho, C. et al., 2009. BLAST+: architecture and applications. BMC bioinformatics, 10(1), p.421. Available at: 10.1186/1471-2105-10-421.

Chang, S. et al., 2011. Mitochondrial genome sequencing helps show the evolutionary mechanism of mitochondrial genome formation in Brassica. BMC genomics, 12(1), p.497. Available at: https://bmcgenomics.biomedcentral.com/articles/10.1186/1471-2164-12-497.

Chang, S. et al., 2013. The mitochondrial genome of soybean reveals complex genome structures and gene evolution at intercellular and phylogenetic levels. PloS one, 8(2), p.e56502. Available at: https://journals.plos.org/plosone/article?id=10.1371/journal.pone.0056502.

Chen, N.W.G. et al., 2018. Common bean subtelomeres are hot spots of recombination and favor resistance gene evolution. Frontiers in plant science, 9, p.1185. Available at: https://www.frontiersin.org/journals/plant-science/articles/10.3389/fpls.2018.01185/full.

Chen, Q. et al., 2021. Harnessing knowledge from maize and rice domestication for new crop breeding. Molecular plant, 14(1), pp.9–26. Available at: https://www.sciencedirect.com/science/article/pii/S1674205220304378#:∼:text=Like%20maize%2C%20cultivated%20rice%20also,more%20amenable%20to%20human%20cultivation.

Choi, I.-S. et al., 2022. Highly Resolved Papilionoid Legume Phylogeny Based on Plastid Phylogenomics. Frontiers in plant science, 13, p.823190. Available at: 10.3389/fpls.2022.823190.

Choi, I.-S. et al., 2021. In and out: Evolution of viral sequences in the mitochondrial genomes of legumes (Fabaceae). Molecular phylogenetics and evolution, 163(107236), p.107236. Available at: https://www.sciencedirect.com/science/article/pii/S105579032100169X.

Cui, H. et al., 2021. Comparative analysis of nuclear, chloroplast, and mitochondrial genomes of watermelon and melon provides evidence of gene transfer. Scientific reports, 11(1), p.1595. Available at: https://www.nature.com/articles/s41598-020-80149-9.

Daniell, H. & Chase, C.D., 2007. Molecular Biology and Biotechnology of Plant Organelles: Chloroplasts and Mitochondria, Springer Science & Business Media. Available at: https://play.google.com/store/books/details?id=-QTmHsc1nRgC.

Darzentas, N., 2010. Circoletto: visualizing sequence similarity with Circos. Bioinformatics (Oxford, England), 26(20), pp.2620–2621. Available at: 10.1093/bioinformatics/btq484.

Day, D., Harvey Millar, A. & Whelan, J., 2013. Plant Mitochondria: From Genome to Function, Springer Science & Business Media. Available at: https://play.google.com/store/books/details?id=6nfkBwAAQBAJ.

Diamond, J., 2002. Evolution, consequences and future of plant and animal domestication. Nature, 418(6898), pp.700–707. Available at: https://www.nature.com/articles/nature01019.

Douce, R., 2012. Mitochondria in Higher Plants: Structure, Function, and Biogenesis, Elsevier. Available at: https://play.google.com/store/books/details?id=bPUX8UckVvsC.

Doyle, J., 1991. DNA Protocols for Plants. In Molecular Techniques in Taxonomy. Berlin, Heidelberg: Springer Berlin Heidelberg, pp. 283–293. Available at: https://link.springer.com/chapter/10.1007/978-3-642-83962-7_18.

Du, L. et al., 2018. Krait: an ultrafast tool for genome-wide survey of microsatellites and primer design. Bioinformatics (Oxford, England), 34(4), pp.681–683. Available at: https://academic.oup.com/bioinformatics/article/34/4/681/4557187.

Felsted, R.L. et al., 1977. Recombinations of subunits of Phaseolus vulgaris isolectins. The journal of biological chemistry, 252(9), pp.2967–2971. Available at: https://www.sciencedirect.com/science/article/pii/S0021925817404571.

Feng, X. & Xu, L., 2018. Mitochondrial genome and environmental adaptation in legumes: insights from tropical species. Frontiers in Plant Science, 9.

Gao, B. et al., 2020. Phylogenomic synteny network analyses reveal ancestral transpositions of auxin response factor genes in plants. Plant methods, 16(1), p.70. Available at: https://plantmethods.biomedcentral.com/articles/10.1186/s13007-020-00609-1.

Greiner, S., Lehwark, P. & Bock, R., 2019. OrganellarGenomeDRAW (OGDRAW) version 1.3.1: expanded toolkit for the graphical visualization of organellar genomes. Nucleic acids research, 47(W1), pp.W59–W64. Available at: 10.1093/nar/gkz238.

Gualberto, J.M. & Newton, K.J., 2017. Plant Mitochondrial Genomes: Dynamics and Mechanisms of Mutation. Annual review of plant biology, 68, pp.225–252. Available at: 10.1146/annurev-arplant-043015-112232.

Guy, L., Kultima, J.R. & Andersson, S.G.E., 2010. genoPlotR: comparative gene and genome visualization in R. Bioinformatics (Oxford, England), 26(18), pp.2334–2335. Available at: 10.1093/bioinformatics/btq413.

Ho, W.K. et al., 2024. A genomic toolkit for winged bean Psophocarpus tetragonolobus. Nature communications, 15(1), p.1901. Available at: https://www.nature.com/articles/s41467-024-45048-x.

Ilic, K., SanMiguel, P.J. & Bennetzen, J.L., 2003. A complex history of rearrangement in an orthologous region of the maize, sorghum, and rice genomes. Proceedings of the National Academy of Sciences of the United States of America, 100(21), pp.12265–12270. Available at: https://www.pnas.org/doi/10.1073/pnas.1434476100.

Indrabalan, U.B. et al., 2021. An extensive evaluation of codon usage pattern and bias of structural proteins p30, p54 and, p72 of the African swine fever virus (ASFV). Virusdisease, 32(4), pp.810–822. Available at: https://pmc.ncbi.nlm.nih.gov/articles/PMC8630154/.

Iriarte, A., Lamolle, G. & Musto, H., 2021. Codon Usage Bias: An Endless Tale. Journal of molecular evolution, 89(9-10), pp.589–593. Available at: 10.1007/s00239-021-10027-z.

Jia, W. & Higgs, P.G., 2008. Codon usage in mitochondrial genomes: distinguishing context-dependent mutation from translational selection. Molecular biology and evolution, 25(2), pp.339–351. Available at: https://academic.oup.com/mbe/article/25/2/339/1132544.

Jin, J.-J. et al., 2020. GetOrganelle: a fast and versatile toolkit for accurate de novo assembly of organelle genomes. Genome biology, 21(1), p.241. Available at: https://genomebiology.biomedcentral.com/articles/10.1186/s13059-020-02154-5.

Katoh, K. et al., 2002. MAFFT: a novel method for rapid multiple sequence alignment based on fast Fourier transform. Nucleic acids research, 30(14), pp.3059–3066. Available at: 10.1093/nar/gkf436.

Kimura, M., 1981. Possibility of extensive neutral evolution under stabilizing selection with special reference to nonrandom usage of synonymous codons. Proceedings of the National Academy of Sciences of the United States of America, 78(9), pp.5773–5777. Available at: 10.1073/pnas.78.9.5773.

Kovar, L. et al., 2018. Gene expression and recombination activity shaped by a distinct mitochondrial genome architecture in Medicago truncatula. Plant Cell, 30(8), pp.1888–1904. Available at: 10.1105/tpc.18.00157.

Kozlov, A.M. et al., 2019. RAxML-NG: a fast, scalable and user-friendly tool for maximum likelihood phylogenetic inference. Bioinformatics (Oxford, England), 35(21), pp.4453–4455. Available at: https://academic.oup.com/bioinformatics/article/35/21/4453/5487384.

Krishnamurthy, P. et al., 2022. Phylogenomic classification and synteny network analyses deciphered the evolutionary landscape of aldo-keto reductase (AKR) gene superfamily in the plant kingdom. Gene, 816(146169), p.146169. Available at: https://www.sciencedirect.com/science/article/abs/pii/S0378111921007642.

Kubo, T. & Mikami, T., 2007. Organization and variation of angiosperm mitochondrial genome. Physiologia plantarum, 129(1), pp.6–13. Available at: 10.1111/j.1399-3054.2006.00768.x.

Kubo, T. & Newton, K.J., 2008. Angiosperm mitochondrial genomes and mutations. Mitochondrion, 8(1), pp.5–14. Available at: 10.1016/j.mito.2007.10.006.

Kubo, T. & Newton, K.J., 2008. Plant mitochondrial genomes and their recombination behaviors. Annual Review of Plant Biology, 59, pp.561–575.

Kühn, K. & Gualberto, J.M., 2012. Recombination in the stability, repair and evolution of the mitochondrial genome. In L. Maréchal-Drouard, ed. Advances in Botanical Research. Advances in botanical research. Elsevier, pp. 215–252. Available at: https://www.sciencedirect.com/science/article/abs/pii/B9780123942791000090.

Kumar, S. & Singh, R., 2020. Mitochondrial genomics in leguminous crops: Adaptation to abiotic stress. Journal of Plant Physiology, 246.

Kurtz, S. et al., 2001. REPuter: the manifold applications of repeat analysis on a genomic scale. Nucleic acids research, 29(22), pp.4633–4642. Available at: 10.1093/nar/29.22.4633.

Lee, C. et al., 2021. The chicken or the egg? Plastome evolution and an independent loss of the inverted repeat in papilionoid legumes. The Plant journal: for cell and molecular biology, 107(3), pp.861–875. Available at: https://onlinelibrary.wiley.com/doi/abs/10.1111/tpj.15351.

Lepcha, P. et al., 2017. A review on current status and future prospects of winged bean (Psophocarpus tetragonolobus) in tropical agriculture. Plant foods for human nutrition (Dordrecht, Netherlands), 72(3), pp.225–235. Available at: https://idp.springer.com/authorize/casa?redirect_uri=https://link.springer.com/article/10.1007/s11130-017-0627-0&casa_token=KZlQ0EEGdJUAAAAA:Va8UF7Dh7czaFhxIXcZAOdtWZ0Ai1g8eUjkx2EtGHyBoL-eeXAX2cNB5dPTJMEaLyLq2d47pZysHPnQ.

Li, G. et al., 2024. Comparative analysis of chloroplast and mitochondrial genomes of sweet potato provides evidence of gene transfer. Scientific reports, 14(1), p.4547. Available at: https://pmc.ncbi.nlm.nih.gov/articles/PMC10894244/.

Li, Y. et al., 2023. An analysis of codon utilization patterns in the chloroplast genomes of three species of Coffea. BMC genomic data, 24(1), p.42. Available at: https://bmcgenomdata.biomedcentral.com/articles/10.1186/s12863-023-01143-4#:∼:text=PR2%2Dplot,selection%20and%20other%20unknown%20elements.

Lu, G. et al., 2023. Assembly and analysis of the first complete mitochondrial genome of Punica granatum and the gene transfer from chloroplast genome. Frontiers in plant science, 14, p.1132551. Available at: https://www.frontiersin.org/journals/plant-science/articles/10.3389/fpls.2023.1132551/full.

Manchekar, M. et al., 2006. DNA recombination activity in soybean mitochondria. Journal of molecular biology, 356(2), pp.288–299. Available at: https://pubmed.ncbi.nlm.nih.gov/16376379/.

Maréchal, A. & Brisson, N., 2010. Recombination and the maintenance of plant organellar genome stability. New Phytologist, 186(2), pp.299–317.

Maréchal, A. & Brisson, N., 2010. Recombination and the maintenance of plant organelle genome stability. The New phytologist, 186(2), pp.299–317. Available at: 10.1111/j.1469-8137.2010.03195.x.

Merritt, B.J. et al., 2015. An empirical review: Characteristics of plant microsatellite markers that confer higher levels of genetic variation. Applications in plant sciences, 3(8), p.1500025. Available at: https://pmc.ncbi.nlm.nih.gov/articles/PMC4542939/.

Mirarab, S. et al., 2014. ASTRAL: genome-scale coalescent-based species tree estimation. Bioinformatics (Oxford, England), 30(17), pp.i541–8. Available at: https://academic.oup.com/bioinformatics/article/30/17/i541/200803.

Morton, B.R. & Wright, S.I., 2007. Selective constraints on codon usage of nuclear genes from Arabidopsis thaliana. Molecular biology and evolution, 24(1), pp.122–129. Available at: 10.1093/molbev/msl139.

Mower, J.P., Sloan, D.B. & Alverson, A.J., 2012. Plant Mitochondrial Genome Diversity: The Genomics Revolution. In Plant Genome Diversity Volume 1. Vienna: Springer Vienna, pp. 123–144. Available at: https://link.springer.com/chapter/10.1007/978-3-7091-1130-7_9.

Murthazar Naim, R., 2020. Growth, physiological and biochemical responses of winged bean (Psophocarpus tetragonolobus) towards different shade levels/Murthazar Naim Raai. Available at: http://studentsrepo.um.edu.my/id/eprint/12372.

Nakamura, T. et al., 2018. Parallelization of MAFFT for large-scale multiple sequence alignments. Bioinformatics (Oxford, England), 34(14), pp.2490–2492. Available at: 10.1093/bioinformatics/bty121.

Niu, Y., Gao, C. & Liu, J., 2022. Complete mitochondrial genomes of three Mangifera species, their genomic structure and gene transfer from chloroplast genomes. BMC genomics, 23(1), p.147. Available at: https://link.springer.com/article/10.1186/s12864-022-08383-1.

Nwokolo, E., 1996. Winged bean (Psophocarpus tetragonolobus(L.) DC.). In Food and Feed from Legumes and Oilseeds. Boston, MA: Springer US, pp. 173–181. Available at: 10.1007/978-1-4613-0433-3_17.

Park, H.-S. et al., 2021. Inheritance of chloroplast and mitochondrial genomes in cucumber revealed by four reciprocal F1 hybrid combinations. Scientific reports, 11(1), p.2506. Available at: https://www.nature.com/articles/s41598-021-81988-w.

Parvathy, S.T., Udayasuriyan, V. & Bhadana, V., 2022. Codon usage bias. Molecular biology reports, 49(1), pp.539–565. Available at: 10.1007/s11033-021-06749-4.

Paul, P., Malakar, A.K. & Chakraborty, S., 2018. Codon usage vis-a-vis start and stop codon context analysis of three dicot species. Journal of genetics, 97(1), pp.97–107. Available at: https://www.ncbi.nlm.nih.gov/pubmed/29666329.

Qiu, S. et al., 2011. Reduced efficacy of natural selection on codon usage bias in selfing Arabidopsis and Capsella species. Genome biology and evolution, 3, pp.868–880. Available at: 10.1093/gbe/evr085.

Rakvong, T. et al., 2024. Tuber development and tuber yield potential of winged bean (Psophocarpus tetragonolobus (L.) DC.), an alternative crop for animal feed. Agronomy (Basel, Switzerland). Available at: https://www.mdpi.com/2073-4395/14/7/1433.

Rao, A. et al., 2024. Codon usage bias in the chloroplast genomes of Cymbidium species in Guizhou, China. Suid-Afrikaanse tydskrif vir plantkunde [South African journal of botany], 164, pp.429–437. Available at: https://www.sciencedirect.com/science/article/pii/S0254629923007470#:∼:text=below%20the%20curve).-,2.2.,direction%20of%20the%20base%20shift.

Satrio, R.D. et al., 2023. A complete chloroplast and mitochondrial genome for velvet bean (Mucuna pruriens, Fabaceae), with genome structure and intergenomic sequence transfers analyses. Available at: https://www.researchsquare.com/article/rs-3612837/latest.

Shah, N. et al., 2020. Extreme genetic signatures of local adaptation during Lotus japonicus colonization of Japan. Nature communications, 11(1), p.253. Available at: https://www.nature.com/articles/s41467-019-14213-y.

Shi N, Yuan Y, Huang R, Wen G., 2024. Analysis of codon usage patterns in complete plastomes of four medicinal *Polygonatum* species (Asparagaceae). Front Genet. 12(1), p.3498. Available at: 10.3389/fgene.2024.1401013.

Singh, N.K., Singh, B.K., Giddhi, A., Srivast, H., et al., 2024. Chloroplast genome sequencing in winged bean (Psophocarpus tetragonolobus L.) and comparative analysis with other legumes. Research Square. Available at: https://assets-eu.researchsquare.com/files/rs-4615004/v1/72008313-a654-45c8-b86b-df0f27d10c6d.pdf?c=1721392366.

Singh, N.K., Singh, B.K., Giddhi, A., Srivastava, H., et al., 2024. Chloroplast genome sequencing in winged bean (Psophocarpus tetragonolobus L.) and comparative analysis with other legumes. Available at: https://scholar.google.com/citations?view_op=view_citation&hl=en&citation_for_view=y8cSxGEAAAAJ:4DMP91E08xMC.

Sinha, K. et al., 2024. Selection on synonymous codon usage in soybean (Glycine max) WRKY genes. Scientific reports, 14(1), p.26530. Available at: https://pubmed.ncbi.nlm.nih.gov/39489740/.

Skippington, E. et al., 2015. Miniaturized mitogenome of the parasitic plant Viscum scurruloideum is extremely divergent and dynamic and has lost all nad genes. Proceedings of the National Academy of Sciences of the United States of America, 112(27), pp.E3515–24. Available at: 10.1073/pnas.1504491112.

Sloan, D.B. et al., 2012. Rapid evolution of enormous, multichromosomal genomes in flowering plant mitochondria with exceptionally high mutation rates. PLoS biology, 10(1), p.e1001241. Available at: https://journals.plos.org/plosbiology/article?id=10.1371/journal.pbio.1001241.

Sloan, D.B., Havird, J.C. & Sharbrough, J., 2017. The on-again, off-again relationship between mitochondrial genomes and species boundaries. Molecular ecology, 26(8), pp.2212–2236. Available at: 10.1111/mec.13959.

Smith, D.R. & Keeling, P.J., 2015. Mitochondrial and plastid genome architecture: Reoccurring themes, but significant differences at the extremes. Proceedings of the National Academy of Sciences of the United States of America, 112(33), pp.10177–10184. Available at: 10.1073/pnas.1422049112.

Sun, M. et al., 2024. Description of mitochondrial genomes and phylogenetic analysis of Megophthalminae (Hemiptera: Cicadellidae). Journal of insect science, 24(6). Available at: https://pmc.ncbi.nlm.nih.gov/articles/PMC11631095/.

Susanti, D., Melati, M. & Kurniawati, A., 2022. Identification of secondary metabolite compounds in two varieties of young winged beans (Psophocarpus tetragonolobus L.) at two harvest ages. Journal of tropical crop science, 9(01), pp.52–67. Available at: https://core.ac.uk/download/pdf/524549461.pdf.

Tanzi, A.S. et al., 2019. Winged bean (Psophocarpus tetragonolobus (L.) DC.) for food and nutritional security: synthesis of past research and future direction. Planta, 250(3), pp.911–931. Available at: https://link.springer.com/article/10.1007/s00425-019-03141-2.

Tillich, M. et al., 2017. GeSeq - versatile and accurate annotation of organelle genomes. Nucleic acids research, 45(W1), pp.W6–W11. Available at: 10.1093/nar/gkx391.

Uliano-Silva, M. et al., 2023. MitoHiFi: a python pipeline for mitochondrial genome assembly from PacBio high fidelity reads. BMC bioinformatics, 24(1), p.288. Available at: https://bmcbioinformatics.biomedcentral.com/articles/10.1186/s12859-023-05385-y.

Wang, D. et al., 2007. Transfer of chloroplast genomic DNA to mitochondrial genome occurred at least 300 MYA. Molecular biology and evolution, 24(9), pp.2040–2048. Available at: https://academic.oup.com/mbe/article/24/9/2040/2925702.

Wang, L. et al., 2018. Genome-wide analysis of codon usage bias in four sequenced cotton species. PloS one, 13(3), p.e0194372. Available at: 10.1371/journal.pone.0194372.

Wang, Y. et al., 2023. Comparative analysis of codon usage patterns in chloroplast genomes of ten Epimedium species. BMC genomic data, 24(1), p.3. Available at: https://bmcgenomdata.biomedcentral.com/articles/10.1186/s12863-023-01104-x.

Wee, C.-C. et al., 2022. Mitochondrial genome of Garcinia mangostana L. variety Mesta. Scientific reports, 12(1), p.9480. Available at: https://www.nature.com/articles/s41598-022-13706-z.

Wenger, A.M. et al., 2019. Accurate circular consensus long-read sequencing improves variant detection and assembly of a human genome. Nature biotechnology, 37(10), pp.1155–1162. Available at: https://www.nature.com/articles/s41587-019-0217-9.

Wicke, S. et al., 2011. The evolution of the plastid chromosome in land plants: gene content, gene order, gene function. Plant molecular biology, 76(3-5), pp.273–297. Available at: 10.1007/s11103-011-9762-4.

Wick, R.R. et al., 2015. Bandage: interactive visualization of de novo genome assemblies. Bioinformatics (Oxford, England), 31(20), pp.3350–3352. Available at: https://pubmed.ncbi.nlm.nih.gov/26099265/.

Wynn, E.L. & Christensen, A.C., 2019. Repeats of unusual size in plant mitochondrial genomes: Identification, incidence and evolution. G3 (Bethesda, Md.), 9(2), pp.549–559. Available at: https://pmc.ncbi.nlm.nih.gov/articles/PMC6385970/.

Xiao, M. et al., 2024. Comparative analysis of codon usage patterns in the chloroplast genomes of nine forage legumes. Physiology and molecular biology of plants: an international journal of functional plant biology, 30(2), pp.153–166. Available at: https://link.springer.com/article/10.1007/s12298-024-01421-0#:∼:text=PR2%2Dplot,of%20the%20nine%20forage%20legumes.

Yang, S., Li, G. & Li, H., 2023. Molecular characterizations of genes in chloroplast genomes of the genus Arachis L. (Fabaceae) based on the codon usage divergence. PloS one, 18(3), p.e0281843. Available at: 10.1371/journal.pone.0281843.

Zhang, C. et al., 2018. ASTRAL-III: polynomial time species tree reconstruction from partially resolved gene trees. BMC bioinformatics, 19(Suppl 6), p.153. Available at: https://bmcbioinformatics.biomedcentral.com/articles/10.1186/s12859-018-2129-y.

Zhang, T. et al., 2012. The complete chloroplast and mitochondrial genome sequences of Boea hygrometrica: insights into the evolution of plant organellar genomes. PloS one, 7(1), p.e30531. Available at: https://pmc.ncbi.nlm.nih.gov/articles/PMC3264610/.

Zhang, Z., 2022. KaKs_Calculator 3.0: Calculating selective pressure on coding and non-coding sequences. Genomics, proteomics & bioinformatics, 20(3), pp.536–540. Available at: https://pubmed.ncbi.nlm.nih.gov/34990803/.

Zhao, T. et al., 2021. Whole-genome microsynteny-based phylogeny of angiosperms. Nature communications, 12(1), p.3498. Available at: https://www.nature.com/articles/s41467-021-23665-0.

