## Supplementary material for "Assembly, Annotation, and Comparative Analysis of the Mitochondrial Genome of Winged Bean (*Psophocarpus tetragonolobus*): Insights into Evolutionary Adaptation and Codon Usage Bias": Supplental Figures S1

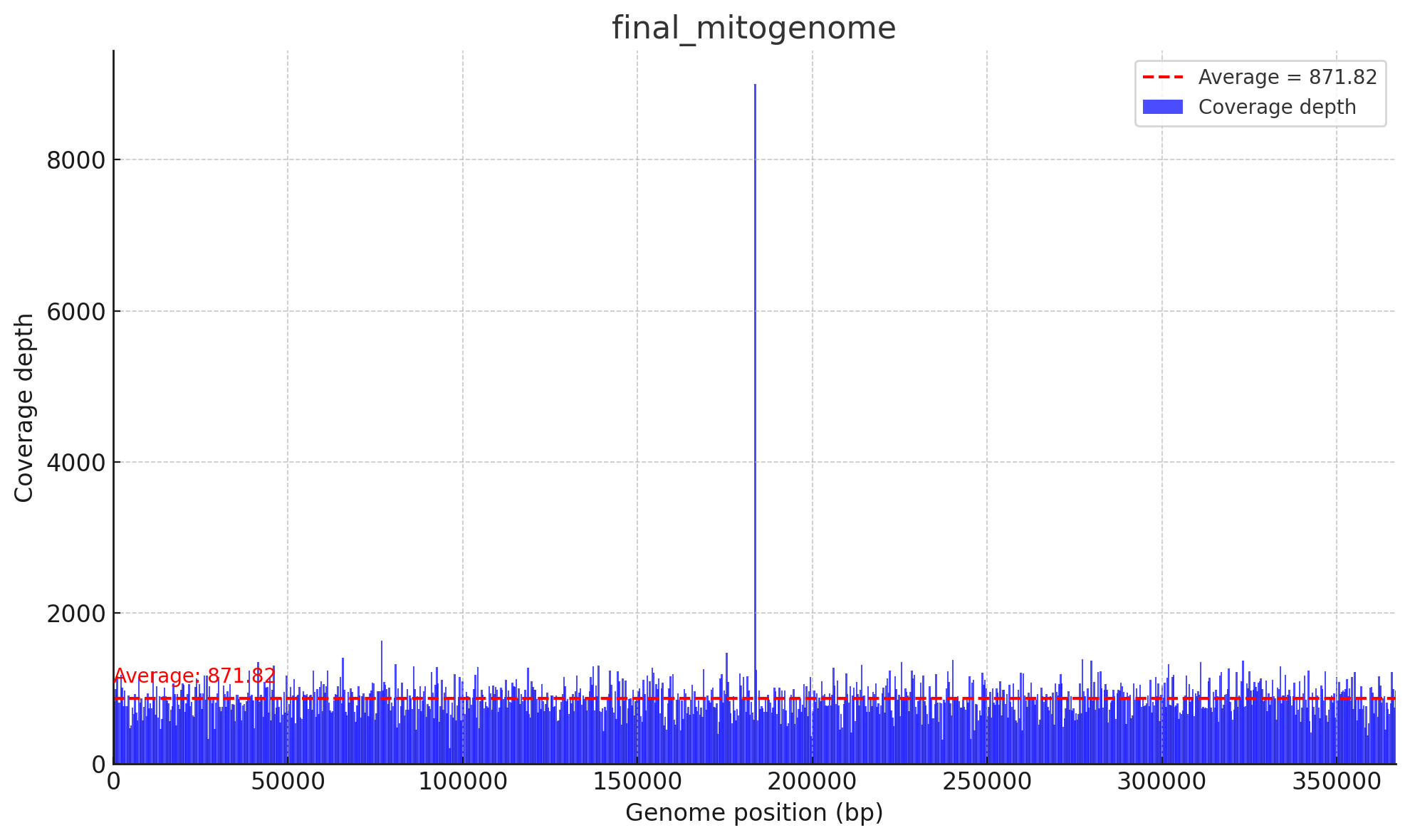

**Supplementary Figure S1:** Coverage depth across genome positions (0–366,925 bp) for the final mitochondrial genome. Blue bars show coverage depth, with a red dashed line marking the average depth (871.82).

Supplementary Figure S2

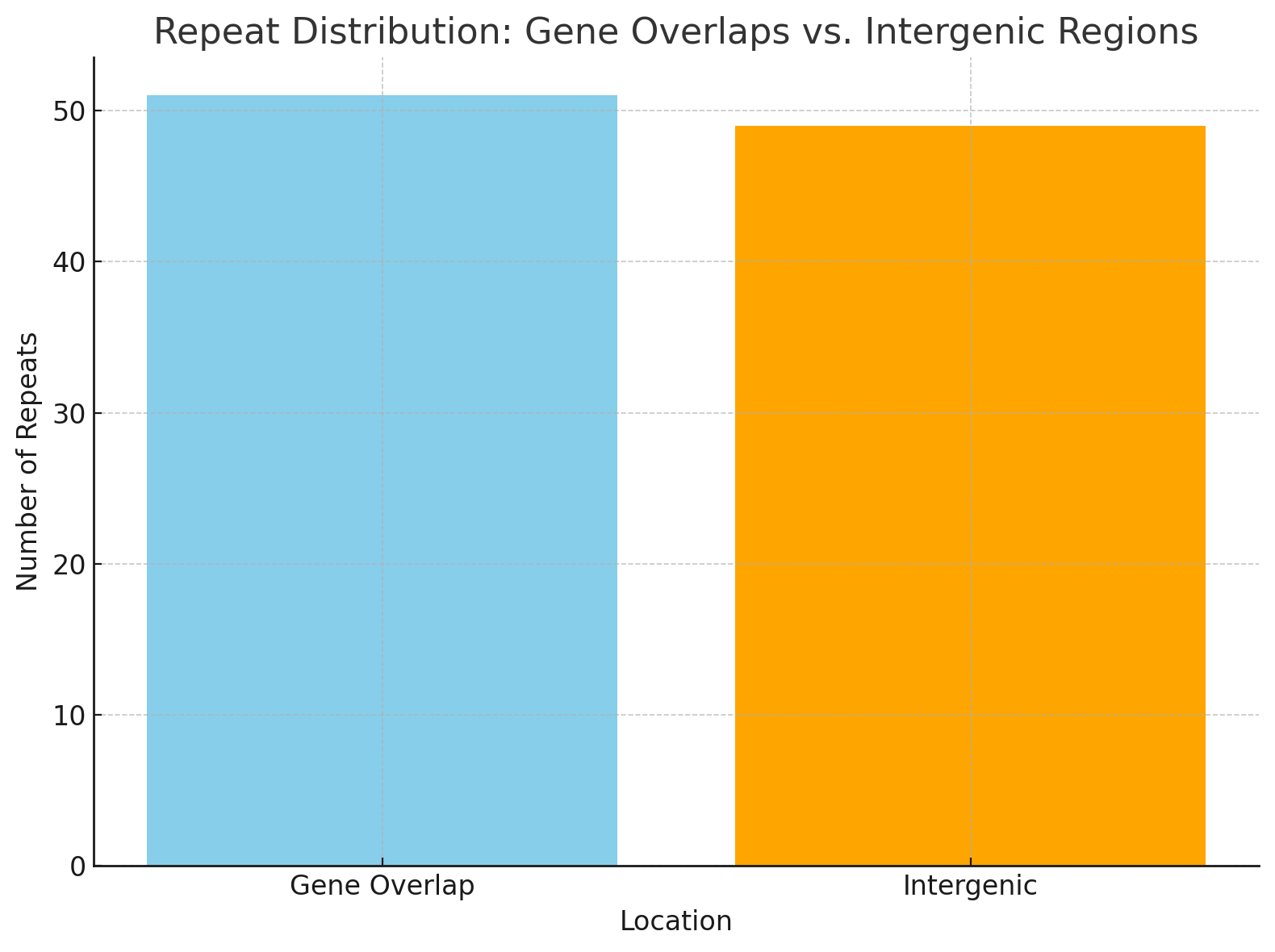

**Supplementary Figure S2: *Repeat content analysis in the Psophocarpus tetragonolobus mitochondrial genome*.** The diagram illustrates the distribution and classification of repeats, including forward, palindromic, and tandem repeats. Repeat types are categorized based on their length and genomic location, highlighting conserved patterns and unique features within the mitochondrial genome.

Supplementary Figure S3

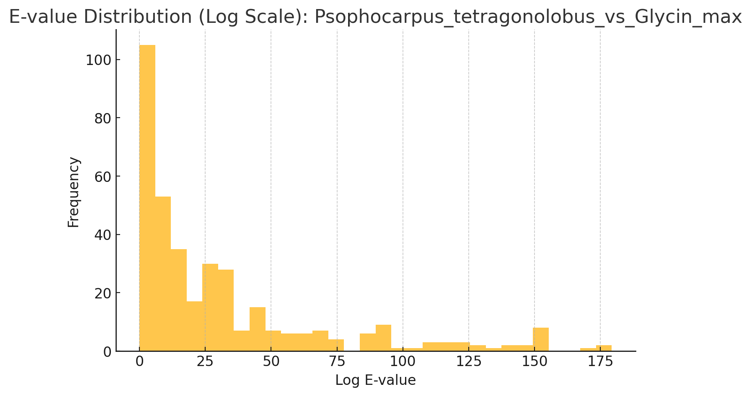

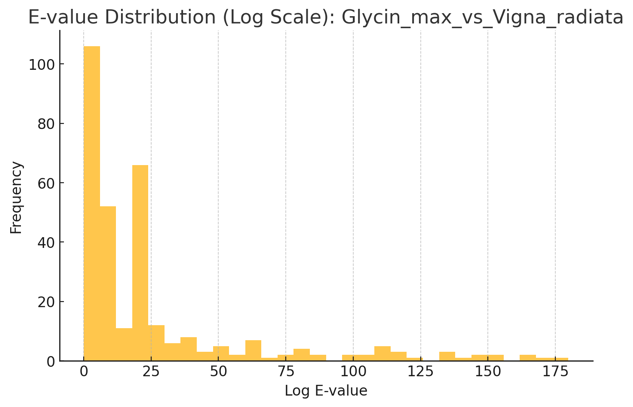

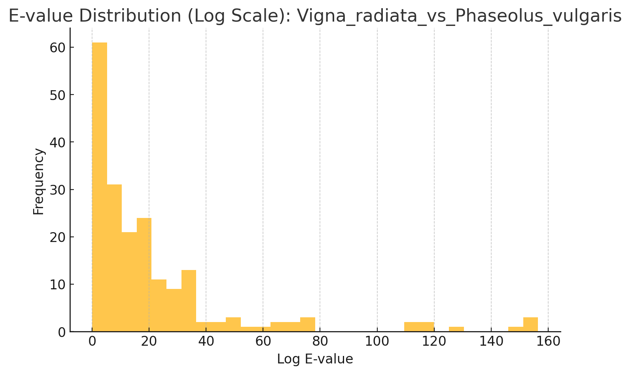

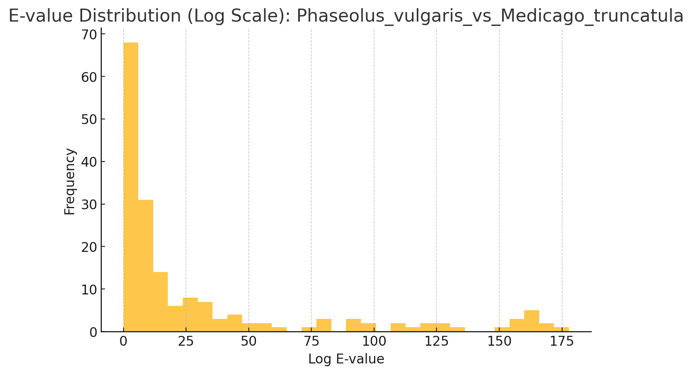

**Supplementary Figure S3:** Distribution of E value (log scale) for each pairwise BLAST performed for synteny analysis
